# Visual Artificial Grammar Learning Across One Year in 7-Year-Olds and Adults

**DOI:** 10.1101/2023.09.13.557366

**Authors:** Daniela K. Schönberger, Patrick Bruns, Brigitte Röder

## Abstract

Acquiring sequential information is of utmost importance, e.g., for language acquisition in children. Yet, the long-term storage of statistical learning in children is poorly understood. To address this question, 27 seven-year-olds and 28 young adults completed four sessions of visual sequence learning (Year 1). From this sample, 16 seven-year-olds and 20 young adults participated in another four equivalent sessions after a 12-month-delay (Year 2). The first three sessions of each year used stimulus set-1, while the last session used stimulus set-2 to investigate transfer effects. Each session consisted of alternating learning and test phases in a modified artificial grammar learning task. In Year 1, seven-year-olds and adults learned the regularities and showed transfer to stimulus set-2. Both groups retained their final performance level over the one-year-period. In Year 2, children and adults continued to improve with stimulus set-1, but did not show additional transfer gains. Adults overall outperformed children, but transfer effects were indistinguishable between both groups. The present results suggest that long-term memory traces are formed from repeated sequence learning which can be used to generalize sequence rules to new visual input. However, the present study did not provide evidence for a childhood advantage in learning and remembering sequence rules.

## Introduction

Both children and adults are capable of tracking sequential information in the environment (Conway, 2020). Statistical learning has been proposed to be particularly effective in children, allowing them, for example, to quickly and implicitly acquire language (Aslin, 2017; Romberg & Saffran, 2010). This assumption has mainly been based on cross-sectional evidence which compared how children vs. adults acquire statistical regularities within single learning sessions: Firstly, behavioral results have suggested a higher sensitivity for visual regularities in young children (< 12 years) compared to older age groups in an implicit learning task (Janacsek et al., 2012). Secondly, neural evidence on implicit learning markers (event-related brain potentials) implied that infants and young children, but not adults, pick up statistical regularities in passive exposure situations, that is, when they are not task-relevant (Mueller et al., 2018; Rohlf et al., 2017). Both lines of evidence were interpreted as a transition from more implicit to more explicit learning mechanisms across development (Daltrozzo & Conway, 2014; Nemeth et al., 2013). It has been hypothesized that this developmental shift underlies children’s superiority in grammar learning or word segmentation (Finn et al., 2014; Smalle et al., 2022). In fact, promoting explicit learning strategies by task instructions (“effortful” learning) impaired grammar learning as compared to passive listening still in adults (Finn et al., 2014). In the same vein, inducing child-like learning mechanisms by decreasing cognitive control prior to passive exposure to hidden auditory regularities, fostered implicit word learning in adults (Smalle et al., 2022). Preventing interference from explicit learning strategies might consequently help to mitigate the age-related sensitivity loss towards passively encountered regularities. The proposal of a high initial sensitivity for statistical regularities which decreases later in childhood has, however, not remained uncriticized.

Some authors have even postulated an improvement in implicit learning of sequential and probabilistic relationships from childhood to adulthood (see, e.g., Lukács & Kemény, 2015). Using transcranial direct current stimulation to disrupt the activity of the dorsolateral prefrontal cortex, which is known to contribute to explicit learning in adults during the acquisition of statistical regularities, caused adults to engage more implicit learning mechanisms similar to those typically shown by children (Friederici et al., 2013). When in adults the same brain area was deactivated during language exposure, word-form learning (Smalle, Panouilleres, et al., 2017) and word segmentation performance were found to be enhanced (Smalle et al., 2022). Following the disruption of dorsolateral prefrontal cortex activity during a first learning phase, delayed learning of visuomotor regularities was reported to be improved when assessed after a 24-hour consolidation period (Ambrus et al., 2020). This finding suggests that implicit learning might improve retention, too. However, longitudinal designs which directly compare multi-session learning and retention effects between children and adults are rare (but see Ferman & Karni, 2010; Smalle, Page, et al., 2017). Thus, it still remains unknown whether children’s learning of rules is more efficient and effective than in adults.

Previous multi-session learning studies with auditory sequences investigated phonological word-form learning (Smalle, Page, et al., 2017) and the learning of an artificial morphological rule (Ferman & Karni, 2010). These studies provided first evidence that a repeated training with a single sequence rule over multiple sessions improves performance in adults and in 8- to 12-year old children alike. Nevertheless, these authors found evidence that learning relied more on explicit knowledge of sequence rules in adults than in children. Results regarding age differences in the retention of implicitly learned sequences were inconsistent: Smalle, Page, et al. (2017) observed memory advantages in 8 to 9 year-old children compared to adults for retaining an implicitly acquired syllable sequence up to 12 months after their last learning session. For retention after a 12-month delay, this age effect only reached significance for matched subgroups with comparable performance levels before the delay. In contrast to implicit learning, an explicitly cued sequence was retained equally well by children and adults across 4 hours, 1 week and 12 months (Smalle, Page, et al., 2017). Ferman and Karni (2010) investigated implicit learning of a sequential language rule across several training sessions and observed preserved performance in 8-year-olds, 12-year-olds and adults after a 2 months retention interval. Recent evidence from an implicit visuomotor task has shown that children aged 9 to 15 years (Tóth-Fáber et al., 2021) and young adults (Kóbor et al., 2017) were able to retain both frequency-based (statistical) knowledge and order-based (sequence) knowledge over an interval of 12 months.

Generalization of rule knowledge to new situations is a crucial skill for learning about the environment or acquiring a language. In single-session transfer studies, adults and children aged 3-6 years (Nowak & Baggio, 2017) or 6-9 years (Jung et al., 2020) respectively, were able to generalize learned regularities to new auditory items (Nowak & Baggio, 2017) and to new instances of the same visual category (Jung et al., 2020). Despite successful transfer in both age groups, Jung et al. (2020) reported that only adults but not children applied explicit rule knowledge in a transfer situation with high retrieval demands (triplet completion task on category level). Moreover, adults and 12-year-olds, but not 8-year-olds, generalized a highly practiced language rule to new items at the end of each training session; two months later, the 12-year-olds and adults were still able to apply their relearned rule knowledge to new items without any performance loss as compared to their transfer performance before the delay (Ferman & Karni, 2010). The authors speculated that 8-year-old children lacked transfer effects, because they failed to explicitly discover the underlying language rule. In fact, in a follow-up study, 8-year-old children were able to transfer the acquired rule knowledge to new items after they had been informed about the rule (Ferman & Karni, 2014). The lack of (implicit) learning transfer in younger children, however, contradicts predictions derived from other lines of research: There is abundant evidence for a shift from generalization to specificity during development; for example, overregularization of grammatical patterns such as past tense of verb forms, in early stages of language learning is a well-documented finding (Marcus et al., 1992). Others (Keresztes et al., 2018) have used the terms pattern completion (generalization) and pattern separation (specificity) to indicate this trend in memory development. Thus, this literature would predict a stronger rather than lower tendency to generalize learned regularities to new instances early in development.

The present longitudinal study investigated how children and adults learn *visual* sequences involving *complex rules* over *several sessions* and whether they *generalize* the rule set *to new surface features*. The present approach extends the existing literature in several ways: (1) Mapping sequence learning across multiple sessions *both* before and after a one-year-delay enabled us to compare learning trajectories of children and adults over an extended period of time. After an initial learning period comprising four sessions, we investigated relearning of previously acquired regularities one year later again over four equivalent sessions. This approach is different from testing retention in a single follow-up session as typically implemented in previous studies (Ferman & Karni, 2010; Kóbor et al., 2017; Smalle, Page, et al., 2017; Tóth-Fáber et al., 2021). (2) We tested the transfer of learning to a new visual stimulus set of a different category instead of testing transfer to new items from the trained category (Ferman & Karni, 2010; Jung et al., 2020; Nowak & Baggio, 2017). (3) Instead of a single rule governing syllable sequences in the auditory domain as implemented in previous multi-session studies (Ferman & Karni, 2010; Smalle, Page, et al., 2017), we used an artificial grammar (AG) learning paradigm, that is, a complex rule set governing picture sequences in the visual modality. This allowed us to directly compare long-term visual learning of complex regularities in children and adults (studied separately in both age groups in Kóbor et al., 2017; Tóth-Fáber et al., 2021), which is important for evaluating whether the previously found age differences are specific to auditory language material or apply to the visual domain as well.

We recruited adults and 7-year-old children, because main differences in both sequence learning and transfer were expected for children younger than 8 years as compared to adults (Keresztes et al., 2018; Mueller et al., 2018). We hypothesized that both children and adults learn the sequence rules and transfer this knowledge to a new stimulus set. Moreover, we expected to observe preserved rule knowledge after a one year period. Seven-year-olds were expected to quicker implicitly acquire the AG, to show lager transfer to a new stimulus set and to feature higher retention of the acquired rule set over one year. Finally, we predicted that children predominantly rely on implicit knowledge while adults acquire more explicit knowledge about the underlying sequence rules.

## Methods

### Participants

The study involved 30 healthy children (7 years old ± 2 months at Session 1), and 30 healthy young adults, mostly undergraduate students recruited at university. All participants did not report a history of seeing or hearing impairments nor any neurological disease. They all were native German speakers.

During the course of the study, the data of five participants had to be excluded from the analyses in Year 1. It turned out that one adult was not a native speaker of German, data of one child was lost due to technical issues, and three participants did not adhere to task instructions at home (one child: one missing session, one child and one adult: no night between two sessions). The remaining 27 seven-year-olds (15 female, mean age at Session 1: 7.05 ± 0.07 years, range: 6.91-7.20 years) and 28 adults (18 female, mean age at Session 1: 23.12 ± 3.50 years, range: 18.83-33.54 years) were included for the analyses of the first four sessions in Year 1 (see Appendix A).

From the original sample of 60 participants, 43 participants completed the experiment a second time after approximately 1 year (see *Study Design*). The data of three of the returning 43 participants (7-year-olds: *n* = 22, adults: *n* = 21) for the home follow-up in Year 2 could not be analyzed due to deviations from task instructions in these sessions (two children: mix-up of stimulus set order, one child: 30 days between two sessions). Since the relearning analysis combined data from Year 1 and Year 2, we additionally had to exclude four of the returning participants since they had missing data in Year 1 for these analyses. This left a total of 16 seven-year-olds (12 female, mean age at first session of Year 2: 8.18 ± 0.09 years, range: 8.03-8.31 years) and 20 adults (14 female, mean age at first session of Year 2: 23.24 ± 2.48 years, range: 19.94-28.47 years) for the analyses of relearning after the one-year-delay (see Appendix A).

Adult participants were monetarily compensated or earned course credit at the end of each session; children received a small toy at the end of each on-site session and after the last session at home in Year 2. Study-related travel costs were reimbursed. All participants, i.e., children and adults, and children’s legal guardians consented prior to participation (with written consent obtained from adult participants and from children’s legal guardians). The study was approved by the Local Ethics Board and was conducted in accordance with the ethical guidelines of the Declaration of Helsinki (revised form of 2013).

## Design and Procedure

### Study Design

All participants completed a total of four sessions in Year 1 on separate days over the time of approximately one week (see Fig. 1A; mean time span Session 1 to Transfer 1 in the final sample: 7.00 ± 0.73 days in 7-year-olds, 6.75 ± 1.08 days in adults):

- Session 1 (in the lab): After the assessment of working memory, the first learning session with the tablet computer (with stimulus set-1) followed. Next, we measured declarative memory and German grammar skills (see *Memory and Language Skills*). Session 1 lasted 90 to 120 minutes including participant briefing and breaks.
- Session 2-3 (at home): Two more learning sessions took place with stimulus set-1 on the tablet computer (mean time span Session 1 to Session 3 in the final sample: 4.11 ± 1.19 days in 7-year-olds, 4.00 ± 1.59 days in adults).
- Transfer 1 (in the lab): On the tablet computer stimulus set-2 was introduced with the same underlying rule set to test transfer of AG learning. Moreover, at the end of the session, explicit knowledge about sequence orders was assessed with a questionnaire in adults and adapted questions with picture cards in children (see *Explicit Knowledge of Sequence Rules*).

**Figure 1.**
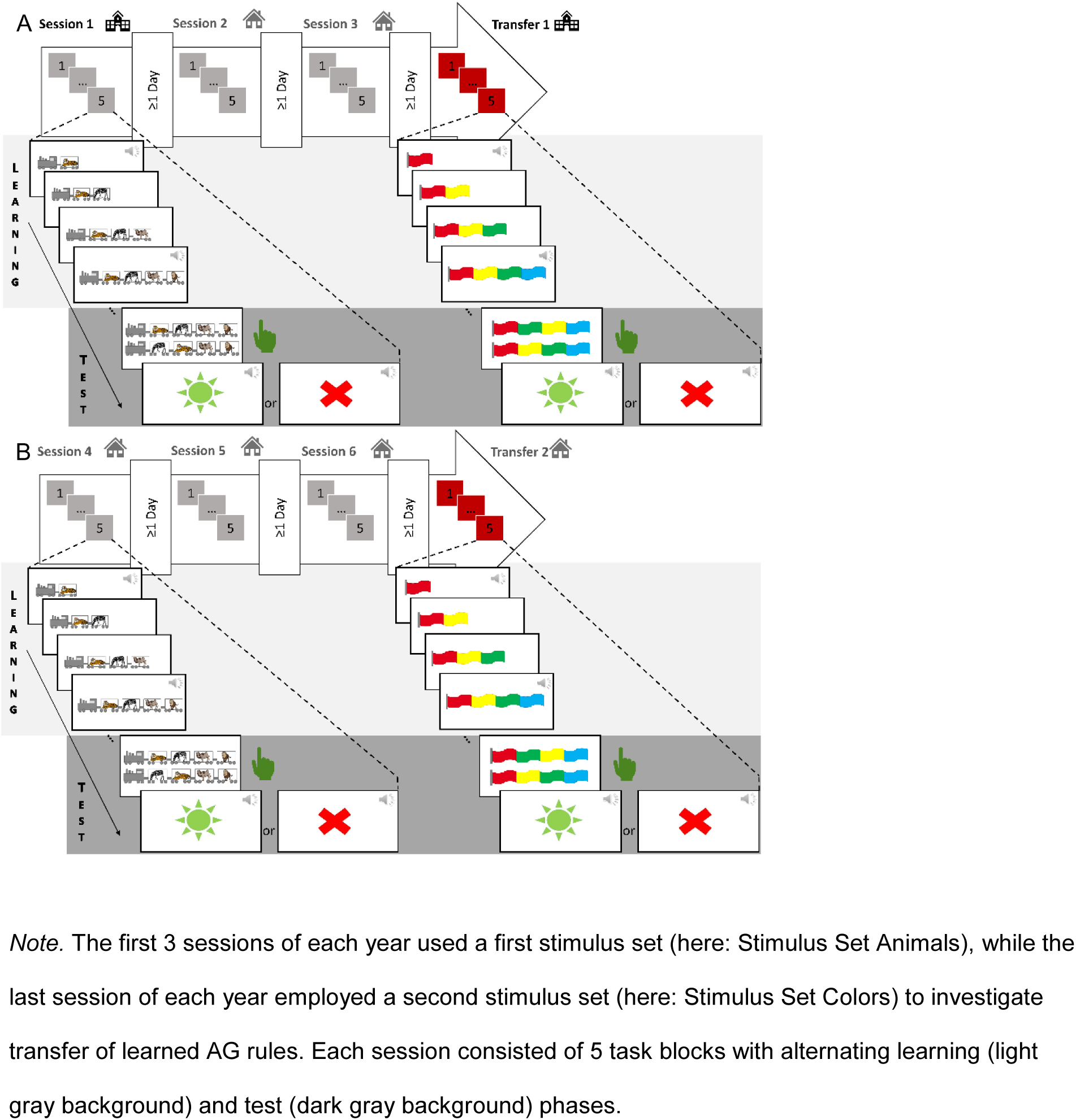
Study Design of all 8 Sessions of Visual Sequence Learning in Year 1 (A) & Year 2 (B)

Due to the ongoing COVID-19 pandemic, the follow-up in Year 2 took place at home. Thus, for these sessions, participants received all materials and a tablet computer by mail and completed four additional learning sessions at home (see Fig. 1B) after approximately one year (mean time span from Session 1 in the final sample: 7-year-olds: 13.00 ± 1.10 months [range: 11.00-15.00 months], adults: 12.75 ± 0.44 [range: 12.00-13.00 months]): Session 4, 5 and 6 used the first stimulus set from Year 1 (that is, the same stimulus set as used in Session 1 to 3), while Transfer 2 employed the second stimulus set from Year 1, that is, the same as used in Transfer 1. All sessions implemented the same AG rule set. At the end of Transfer 2, explicit sequence knowledge was assessed with the same questionnaires as in Year 1 (Transfer 1) which had been mailed to the parents and participants, respectively.

### Visual Sequence Learning Task

The present sequence learning task built on previous AGL tasks for children of similar age (esp. Rosas et al., 2010; Witt & Vinter, 2012). In AGL tasks, participants are exposed in a learning phase to sequences of elements which follow a certain set of sequence rules (a finite state grammar, see Fig. 2). In the following test phase, they have to discriminate grammatical sequences from sequences that violate the sequence rules (“ungrammatical” sequences).

**Figure 2.**
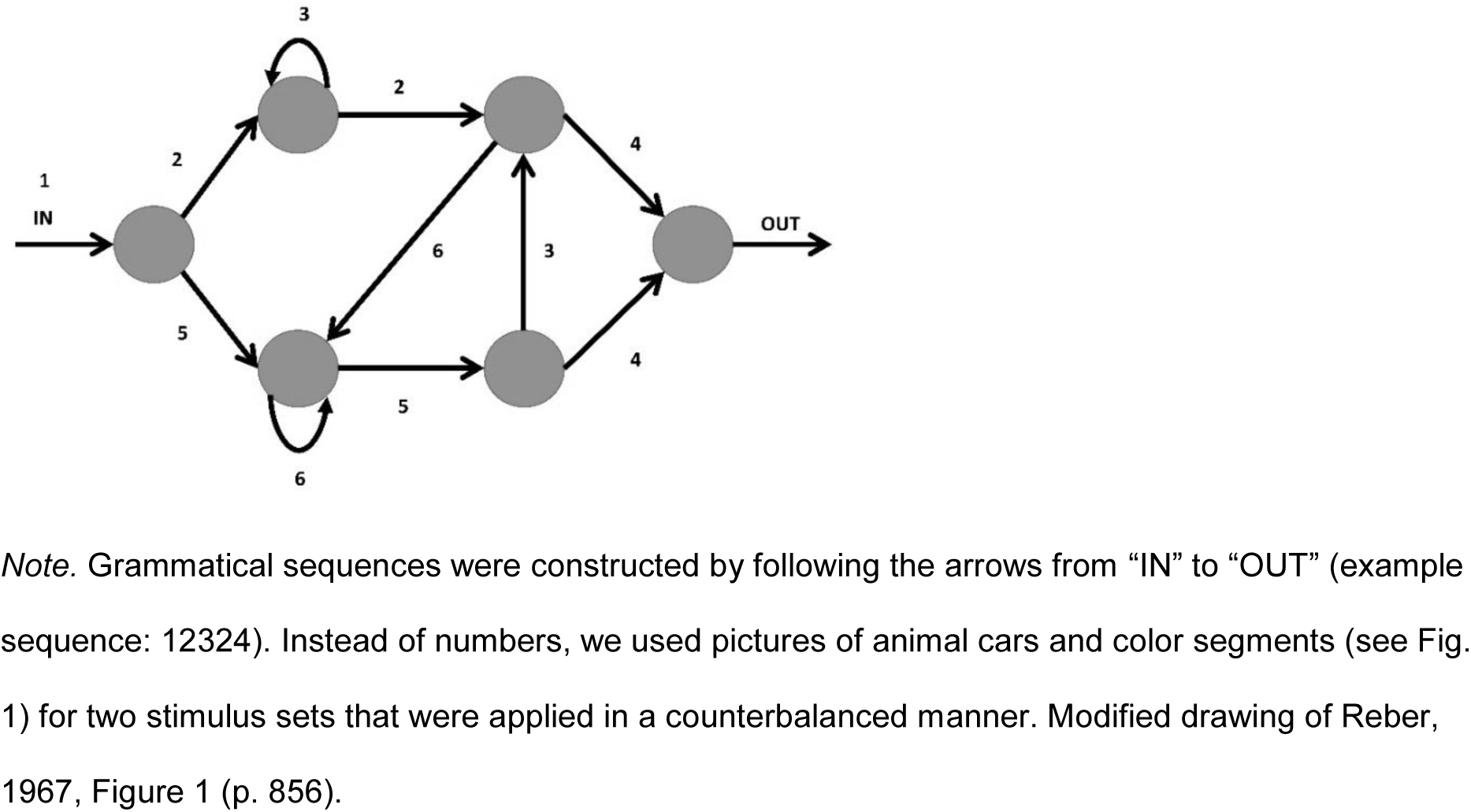
Sequence Rules for the Visual Sequence Learning Task (Artificial Grammar by Reber, 1967)

We implemented visual sequence learning on a tablet computer with five blocks of alternating learning phases (passive watching) and test phases (two-alternative forced choice task) per session. All instructions were child-directed voice recordings embedded in the task and they were automatically played upon starting the application. This procedure guaranteed a high level of standardization of the sessions at home (see Appendix B, Table B.2 for the wording of all instructions).

One task block comprised one learning phase and one subsequent test phase (see Fig. 1). In the learning phases (see Fig. 1 yellow background), participants were instructed to attentively watch 18 grammatical sequences with 3 to 7 items which were randomly taken from the 27 possible grammatical sequences (see *Construction of Grammatical and Ungrammatical Sequences*). Only one stimulus set, either Stimulus Set Animals with animals in train cars (“circus trains” belonging to a circus director: adapted from Rosas et al., 2010) or Stimulus Set Colors with color segments (“team flags” belonging to a sports team: adapted from Witt & Vinter, 2012) was presented. Which stimulus set served as set-1 and set-2 was counterbalanced across participants.

In the test phases following each learning phase (see Fig. 1 green background), participants were asked in each of the 10 trials to select from the two displayed sequences the sequence they considered as belonging to the previously introduced task character (i.e., the grammatical sequence; two-alternative forced choice). This instruction was repeated for each trial of the test phase and participants selected their choice by touching the selected sequence on the tablet’s touchscreen. Each individual test trial comprised two sequences with each being either short (3-5 items, 5 test trials) or long (6-7 items, 5 test trials). Always one of the two sequences was grammatical and one was ungrammatical (see *Construction of Grammatical and Ungrammatical Sequences*). Participants received audio-visual feedback (see *Stimuli and Apparatus*) after each test trial which indicated if their answer was correct or incorrect.

One session of the sequence learning task (5 blocks with an alternating learning and test phase each) took about 25-30 min to complete; additionally, short breaks were offered to the participants after each block.

## Materials

### Stimuli and Apparatus

#### Stimuli and Timing of the Visual Sequence Learning Task

Participants completed the visual sequence learning task in the mobile Neurobs Presentation App (Version 3.0.1, Neurobehavioral Systems Inc., 2019) on a 10.1-inch tablet computer (Samsung Galaxy Tab A 10.1).

Grammatical and ungrammatical sequences were 3 to 7 items long and were built from a total of 5 different pictures per stimulus set (Stimulus Set Animals with circus trains: giraffe, camel, lion, tiger and zebra; Stimulus Set Colors with team flags: blue, yellow, green, purple and red). The individual picture size of 240x240 pixels was 4° of visual angle. Viewing distance was ca. 40 cm, which is used in all subsequent calculations of the visual angle. For grammatical sequences, these pictures were assigned to the positions in the AG (see numbers in Fig. 2) in two different, randomly selected ways per stimulus set to control for position effects of specific animals or colors, respectively (an individual assignment for each participant was technically not feasible due to the implementation in a mobile App). This picture assignment resulted in two task versions available for each stimulus set (see Appendix B, Table B.1), which were used in a counterbalanced manner in both age groups.For the animals of the Stimulus Set Animals, we used pictures from the “Multilingual Picture” database (MultiPic; Dunabeitia et al., 2018). The surrounding train (car) features and all other visual stimuli of the task were custom made digital drawings.

While the recorded instructions played, the screen showed a picture of a scene corresponding to the stimulus set (Stimulus Set Animals: a circus tent; Stimulus Set Colors: a sports stadium; both retrieved from the MultiPic database by Dunabeitia et al., 2018). To start a learning or test phase, participants were asked to click on a green button in the upper half of the screen (size: 4° visual angle).

The duration of a single learning trial varied with the number of items of the presented sequence: items, that is single pictures of animal cars or flag segments, built up sequentially (1 s per item), but the full sequence stayed on the screen for 3 s for all sequence lengths. All learning trials were displayed in the middle of the screen. Each trial started and ended with a sound which was presented synchronously with the first and last item of a sequence. Two different sounds, both retrieved from an open-source website (http://www.findsounds.com, accessed 11/16/18; train whistle for circus trains: http://atsf.railfan.net/airhorns/p5.html/, wind blow for team flags: http://www.anzwad.com/dods/sound/ambient), were used, one for each stimulus set. Each learning phase consisted of 18 learning trials amounting to an average duration of about three minutes.

In the test phase all items of the two sequence were simultaneously displayed from the beginning. Each sequence stayed on the screen until the participant clicked on one of the two sequences which took on average of about two minutes for the 10 test trials in total. For the sequence pairs in each test trial, one sequence was displayed in the upper half of the screen, and the other sequence was displayed in the lower half of the screen, with an equal likelihood for the grammatical sequence of appearing in one of the two screen locations.

For audiovisual feedback, a green sun icon with a corresponding sound (correct) or a red cross icon with a corresponding sound (incorrect) was presented (adapted from Leon Guerrero et al., 2016). Participants all started with the same sound volume and were allowed to customize the volume over the course of the session(s).

After the presentation of each sequence (learning phases) and after the response feedback to each trial (test phases), respectively, a grey fixation cross was shown for 1s in the center of the screen (size: 0.5° visual angle) before the start of the next trial.

#### Construction of Grammatical and Ungrammatical Sequences

We used the AG system introduced by Reber (1967) for constructing grammatical sequences (see Fig. 2). Meta-analyses have attested this AG a relatively low complexity suitable for use in developmental populations (Schiff & Katan, 2014).

From this grammar, 27 grammatical sequences with 3 to 7 items were constructed using the Web App “AGSuite” (Cook et al., 2017; for a full list of all grammatical sequences see Appendix B, Table B.3), which were divided into 9 “short” sequences consisting of 3 to 5 items and 18 “long” sequences consisting of 6 to 7 items. We then compiled a pool of 140 ungrammatical sequences with 3 to 7 items. For sequences of 5 to 7 items this was achieved by randomly shuffling the middle elements of the grammatical sequences (leaving the most salient first and last item unchanged). For grammatical sequences of only 4 items, either the first 3 or the last 3 items were shuffled. This kept the first or the last item, respectively, unchanged to avoid grammaticality judgements predominantly based on these most salient positions. For the two grammatical sequences of only 3 items, the first item stayed the same, while the second and third item swapped places (as all other permutations would result in grammatical sequences). Next we computed a measure called “Global Associative Chunk Strength” (ACS) according to Cook et al. (2017, p. 1648) in a slightly modified way for all the 140 resulting ungrammatical sequences, to match them in difficulty for short (low ACS) and long trials (high ACS): We defined ACS as the sum of an ungrammatical sequence’s shared bigrams (item pairs) and trigrams (item triplets) with all bigrams/trigrams existing in all grammatical sequences of the same length (3 to 5 items or 6 to 7 items, respectively), divided by the total number of bigrams/trigrams in the given ungrammatical sequence: A higher ACS means that an ungrammatical sequence shared more picture pairs and triplets with the grammatical sequences of the same length, making this sequence more difficult to be identified as ungrammatical. From the 30 short ungrammatical sequences (3 to 5 items), 11 sequences with a similar ACS (*M* = 1.44, *SD* =.18; range: 1.25 to 1.75) were selected, which were later randomly paired with the 9 short grammatical sequences to produce the pairs of displayed sequences in the “simple” test trials. For the second trial type, we picked 19 out of the 110 long ungrammatical sequences (6 to 7 items) that had a similar ACS (*M* = 5.97, *SD* =.51; range: 5.03 – 6.85) and later paired them randomly with the 18 long grammatical sequences to make up the “difficult” test trials (for a full list of all ungrammatical sequences see Appendix B, Table B.3). For each test phase, 5 randomly chosen pairs of “simple” sequences (short sequence with a low ACS) and 5 randomly chosen pairs of “difficult” sequences (long sequence with a high ACS) were presented in a random order, adding up to 10 test trials per block.

Each individual grammatical sequence was restricted to appear at most once per learning and once per test phase, and each individual ungrammatical sequence only once per test phase. Thus, within each task block (comprising 1 learning phase and 1 subsequent test phase), an individual ungrammatical sequence was not seen more than once and an individual grammatical sequence was not encountered more than twice.

### Explicit Knowledge of Sequence Rules

Explicit knowledge about underlying rules of the sequence learning task was assessed in the end of Transfer 1 with a questionnaire in adults and adapted questions with picture cards in children, both based on a procedure by Whitmarsh et al. (2013).

The same questionnaires were administered in the end of Transfer 2. Children’s parents were asked to pose the questions and document the answers of their children in Transfer 2, since unlike Transfer 1, this transfer session was run at home. Due to miscommunication, 5 children answered this second questionnaire of Year 2 after Session 6, that is, not to stimulus set-2 used for transfer, but instead to stimulus set-1.

The questionnaire contained 11 (adults) or 5 (children) specific questions, respectively, measuring reported knowledge about legal first and last pictures (*With what [animal/color] could the trains/flags of [introduced person/team] begin/end?*) and legal bigram transitions (*What animals/colors could repeat themselves? What animal(s)/color(s) could follow animal/color X? What animal(s)/color(s) could **not** follow animal/color X?*) as described in detail in Whitmarsh et al. (2013). Adults and children in Year 2 had to choose from the five possible colored pictures (animals/colors) provided as response options in the questionnaire or for children in Year 1 as printed cards (multiple choice).

From these specific questions, we calculated an explicit knowledge score for Year 1 and Year 2, respectively. To this end, correct answers (correctly chosen pictures) were added up and incorrect answers (incorrectly chosen pictures) were subtracted from the correct answers in a weighted manner, in which the answer to each question was weighted according to the probability of valid bigram answers (i.e., the number of correct pictures chosen divided by the number of all possible correct pictures [considering misses] and in an analogue manner for incorrect picture choices [considering correct rejections]), since the number of valid bigram answers differed for different pictures (as they represented different “arrows” in the artificial grammar, see numbers in Fig. 2). This means that for each question, a score from -1 (no correct answers given) to 1 (all correct answers given, without any false alarms) could be obtained. The resulting scores for each question were then added up and divided by the number of questions answered in total, to obtain the final score for explicit sequence knowledge (range: -1 to 1). Higher scores indicated more explicit sequence knowledge.

### Memory and Language Skills

To assess working memory, declarative memory and German grammar skills, we administered equivalent psychometric tests in Session 1 (Year 1) in both age groups and normalized all test scores according to age (except for Plural German Grammar Skills in adults for which norms were not available and for which hence raw scores were analyzed). Table 1 shows the administered subscales of these psychometric tests together with the achieved test scores for both age groups in the final sample of Year 1.

**Table 1.** Subscales for Assessed Memory and Grammar Skills with the Mean (and SD) per Age Group.

| **Cognitive Skill** | **7-Year-Olds (n = 27)** |  | **Adults (n = 28)** |  |
| --- | --- | --- | --- | --- |
| Declarative Memory | Atlantis (Encoding)<sup>1</sup> | 11.67 (2.76) | Rebus (Encoding)<sup>2</sup> | 11.71 (2.17) |
|  | Atlantis Delayed (Retrieval)<sup>1</sup> | 12.37 (2.39) | Rebus Delayed (Retrieval)<sup>2</sup> | 11.86 (1.53) |
| Working Memory | Number Recall<sup>1</sup> | 10.48 (2.62) | Digit Span<sup>3</sup> | 11.11 (2.25) |
| German Grammar I (Plural) | Subscale 8 SET<sup>4</sup> | 57.56 (9.66) | Subtest AR4 D-PA<sup>5</sup> | 12.25 (1.11) |
| German Grammar II (General) | Subscale 9 SET<sup>4</sup> | 60.19 (15.90) | Subscale AG D-PA<sup>5</sup> | 53.46 (9.16) |

*Note.* <sup>1</sup> KABC-II (Melchers & Melchers, 2015): normalized to $M \pm SD$ of $10 \pm 3$ ; <sup>2</sup> K-TIM (Melchers et al., 2006): normalized to $M \pm SD$ of $10 \pm 3$ ; <sup>3</sup> WAIS-IV (Petermann, 2012): normalized to $M \pm SD$ of $10 \pm 3$ ; <sup>4</sup> SET 5-10 (Petermann, 2018): normalized to $M \pm SD$ of $50 \pm 10$ ; <sup>5</sup> D-PA (Rieser & Liepmann, 2014): Subscale AG normalized to $M \pm SD$ of $50 \pm 10$ , Subtest AR4: no age norms available (raw scores).

## Data Analysis

We characterized learning trajectories and averaged performance scores as proportion of correct test trials: Across session learning was assessed as the mean performance of 50 test trials of each session. Within session learning trajectories were derived based on the means of 10 test trials per block.

Trials with shorter reaction times than 200 ms were disregarded, since we did not consider it as feasible to successfully process the two sequences within less than this time. This exclusion criterion reduced trial numbers by 0.18 % (a total of 20 trials from 5 seven-year-olds were excluded) for analyses of Year 1 and by 0.12 % (a total of 17 trials from 7 seven-year-olds and from 1 adult were disregarded) of Year 1 and Year 2 combined for relearning analyses in the returning subsample.

To compare performance changes over time between the two age groups, repeated-measures Analyses of Variance (ANOVAs) were conducted using the *ez* package in R (Lawrence, 2016), with *Age* (7-year-olds vs. adults) as between-subject factor and *Session* (levels depending on analyses as described below and in the respective *Results* section) as within-subject factor.

1. Within Year 1, the following comparisons were analyzed:

- Session 3 vs. Session 1 (Learning Gains)
- Transfer 1 vs. Session 1 (Transfer Savings)
- Transfer 1 vs. Session 3
2. For comparing start and end levels between Year 1 and Year 2, the following sessions were analyzed:

- 4. Session 4 vs. Session 1
- 5. Session 4 vs. Session 3
- 6. Session 6 vs. Session 3
3. For comparing session differences between Year 1 and Year 2 (performance improvement over 3 sessions, transfer performance relative to the first and last session with the first stimulus material), an additional within-subject factor *Year* (Year 1 vs. Year 2) was added in the ANOVAs, resulting in the within-subject factors *Session* and *Year* with levels described in the respective *Results* section.

ANOVAs were followed up with appropriate post-hoc tests. If scores were not normally distributed, Wilcoxon signed rank tests (instead of *t*-tests) and Spearman correlation coefficients (*r_s_*) were calculated. Two tailed significant (<.05) *P*-values (if not indicated otherwise) were Greenhouse-Geisser-corrected (in case of violated sphericity) or Holm-corrected (in case of multiple comparisons). Effect sizes were calculated as generalized eta squared (η^2^_g_) for ANOVAs, as Cohen’s *d* for *t*-tests and as matched rank biserial correlation (*r*) for Wilcoxon singed rank tests, respectively.

For session comparisons that involved Session 1 or Session 4, additional control analyses were conducted with proportion correct of test trials averaged over block 2 to 5 of each session (without the first block), to account for task novelty (see Appendix C, Table C.1-3).

Due to the home-setting, there were some additional deviations from the task instructions in the final sample, mainly in Year 2 (3 participants with 4 instead of 3 sessions with the first stimulus set, 2 participants with 6 instead of 5 task blocks in 1 session, 1 participant with 4 instead of 5 task blocks in Session 5). These participants were included in the final analyses, since they did not show any systematic peculiarities in their response patterns and only the originally scheduled trials were included in case of additional task blocks completed (*n* = 5) or session performance was averaged over the available data in case of a missing task block (*n* = 1), respectively. Additionally, we checked for each of the reported analysis whether excluding these six participants with slightly different task exposure would qualitatively change the pattern of results.

We performed equivalent Bayesian analyses for all inferential statistical analyses in the software JASP (Version 0.14.1; JASP Team, 2021), using default priors, and report the Bayes Factor *(BF_10_).* The *BF* helps evaluating whether the data at hand support the null-hypothesis (H_0_) or the alternative hypothesis (H_1_), and has been described as a suitable tool for interpreting null results (Dienes, 2014). For main and interaction effects in ANOVAs, we report the inclusion Bayes factor *(BF*_incl_) – which compares models that contain the effect of interest to equivalent models stripped of this effect – as implemented in JASP Version 0.14.1 and recommended by e.g. Mathôt (2017). *BF* values between 1/3 and 1/10 indicate moderate evidence for the H_0_, while a BF of lower than 1/10 is considered strong evidence for the H_0_; a BF between 1 and 1/3 is defined as anecdotal evidence for the H_0_ (Schönbrodt & Wagenmakers, 2018). On the other hand, *BF* values between 3 and 10 indicate moderate evidence for the H_1_, while a *BF* from 10 onwards is considered as strong evidence for the H_1_ and a *BF* between 1 and 3 is defined as anecdotal evidence for the H_1_ (Schönbrodt & Wagenmakers, 2018).

All data analyses apart from Bayesian analyses were performed in the software R (Version 4.1.0; R Core Team, 2021).

To further look into within-session learning, we made use of the trial-by-trial response data and fit the state-space random effects model by Smith et al. (2005) to binary responses (correct = 1, incorrect = 0) in all 150 test trials of the 3 sessions with stimulus set-1, separately for each age group and each year (Session 1 to 3 in Year 1: data from 27 children and 28 adults; Session 4 to 6 in Year 2: data from 15 children and 20 adults). This model estimated the first trial as the timepoint where learning had first occurred for the whole population (i.e., age group), by estimating an unobservable learning state process, defined as a random walk. For an estimation of the learning curves, it used a state-space random effects model and Expectation-Maximization algorithm, characterizing the dynamics of the learning process as a function of trial number (Smith et al., 2005). The modeling script was provided in Matlab (Matlab, MathWorks 2020) from the website indicated by Smith et al. (2005; http://annecsmith.net/behaviorallearning.html).The estimated first learning trial from this population modeling was then used to compare within-session learning between age groups.

## Results

### Repeated Learning across one Week (Year 1)

We first tested whether both age groups performed above chance in every session, to assess whether they had learned the AG rule (Fig. 3A). For all sessions, group-level performance exceeded the chance level of 0.5 (two-alternative forced-choice trials) in both 7-year-olds (all *t*(26) ≥ 2.80, all *p* ≤ .005, all *d* ≥ 0.54, all *BF*_+0_ ≥ 9.53; one-sided) and in adults (all *V*(27) = 406.00, all *p* ≤ .004, all *r* ≥ 0.87, all *BF*_+0_ > 100; one-sided). All following analyses are based group *Means(SDs)* in Table 2A (*n =* 27 7- year-olds & *n =* 28 Adults).

**Figure 1a.**
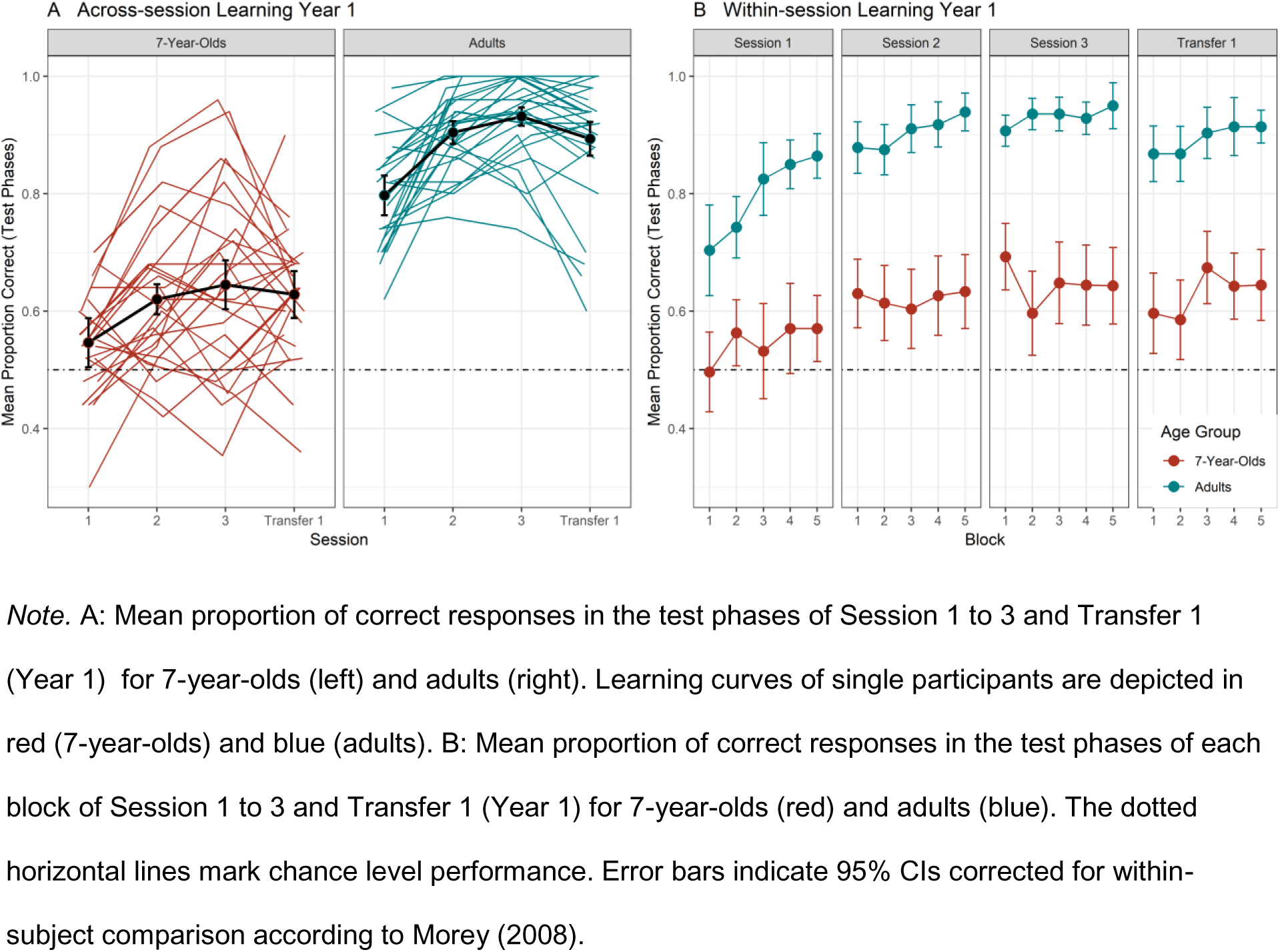
Performance Trajectories Across Sessions and Within Sessions of Year 1.

**Table 2.** A Year 1: Proportion Correct in AGL per Session and Age Group.

A Year 1: Proportion Correct in AGL per Session and Age Group
|  | Session 1 |  | Session 2 |  | Session 3 |  | Transfer 1 |  |
| --- | --- | --- | --- | --- | --- | --- | --- | --- |
|  | 7yo | Ad | 7yo | Ad | 7yo | Ad | 7yo | Ad |
| *N* | 27 | 28 | 27 | 28 | 27 | 28 | 27 | 28 |
| *M* | .55 | .78 | .62 | .90 | .65 | .93 | .63 | .89 |
| *SD* | .09 | .09 | .12 | .07 | .16 | .07 | .11 | .11 |
| Min | .30 | .62 | .42 | .76 | .35 | .74 | .36 | .60 |
| Max | .70 | .98 | .88 | 1.00 | .96 | 1.00 | .90 | 1.00 |
| *N* * | 16 | 20 | 16 | 20 | 16 | 20 | 16 | 20 |
| *M* | .56 | .79 | .69 | .92 | .69 | .94 | .67 | .90 |
| *SD* | .07 | .10 | .13 | .06 | .17 | .07 | .09 | .12 |
| Min | .44 | .62 | .50 | .76 | .46 | .74 | .52 | .60 |
| Max | .70 | .98 | .88 | 1.00 | .96 | 1.00 | .90 | 1.00 |

B Year 2: Proportion Correct in AGL per Session and Age Group
|  | Session 4 |  | Session 5 |  | Session 6 |  | Transfer 2 |  |
| --- | --- | --- | --- | --- | --- | --- | --- | --- |
|  | 7yo | Ad | 7yo | Ad | 7yo | Ad | 7yo | Ad |
| *N* | 16 | 20 | 16 | 20 | 16 | 20 | 16 | 20 |
| *M* | .71 | .91 | .76 | .94 | .80 | .94 | .73 | .92 |
| *SD* | .11 | .10 | .16 | .09 | .13 | .09 | .13 | .09 |
| Min | .52 | .60 | .48 | .70 | .52 | .68 | .49 | .64 |
| Max | .90 | 1.00 | .98 | 1.00 | 1.00 | 1.00 | .98 | 1.00 |

Note. 7yo = 7-year-olds, Ad = Adults, Min = minimal value, Max = maximal value.
\* Subgroup of returning participants with data for Year 1 & Year 2 (see section *Participants*).

### Age Comparison: Performance Improvement and Transfer to new Stimuli

To evaluate benefits of repeated sequence training, we tested to what degree performance improved from the first (Session 1) to the last session (Session 3) with the first stimulus set. To this end, mean session scores (proportion correct of 50 test trials) were entered into an ANOVA with the factors *Age* (between-subject; levels: 7-year-olds, adults) and *Session* (within-subject; levels: Session 1, Session 3): Learning performance improved in both groups from Session 1 to Session 3 (see Table 2; main effect of *Session: F*(1, 53) = 40.59, *p* < .001, η^2^_g_ = .24; *BF*_incl_ > 100), as reflected in the positive mean difference scores depicted in Figure 4A. The mean proportion of correct responses was higher in adults compared to 7-year-olds (see Table 2) for both sessions (main effect of *Age: F*(1, 53) = 152.93, *p* < .001, η^2^_g_ = .63, *BF*_incl_ > 100), while Learning Gains from Session 1 to Session 3 did not significantly differ between groups (interaction *Age*\**Session*: *F*(1, 53) = 0.95, *p* = .333, η^2^_g_ = .01; *BF*_incl_ = 0.40).

**Figure 4.**
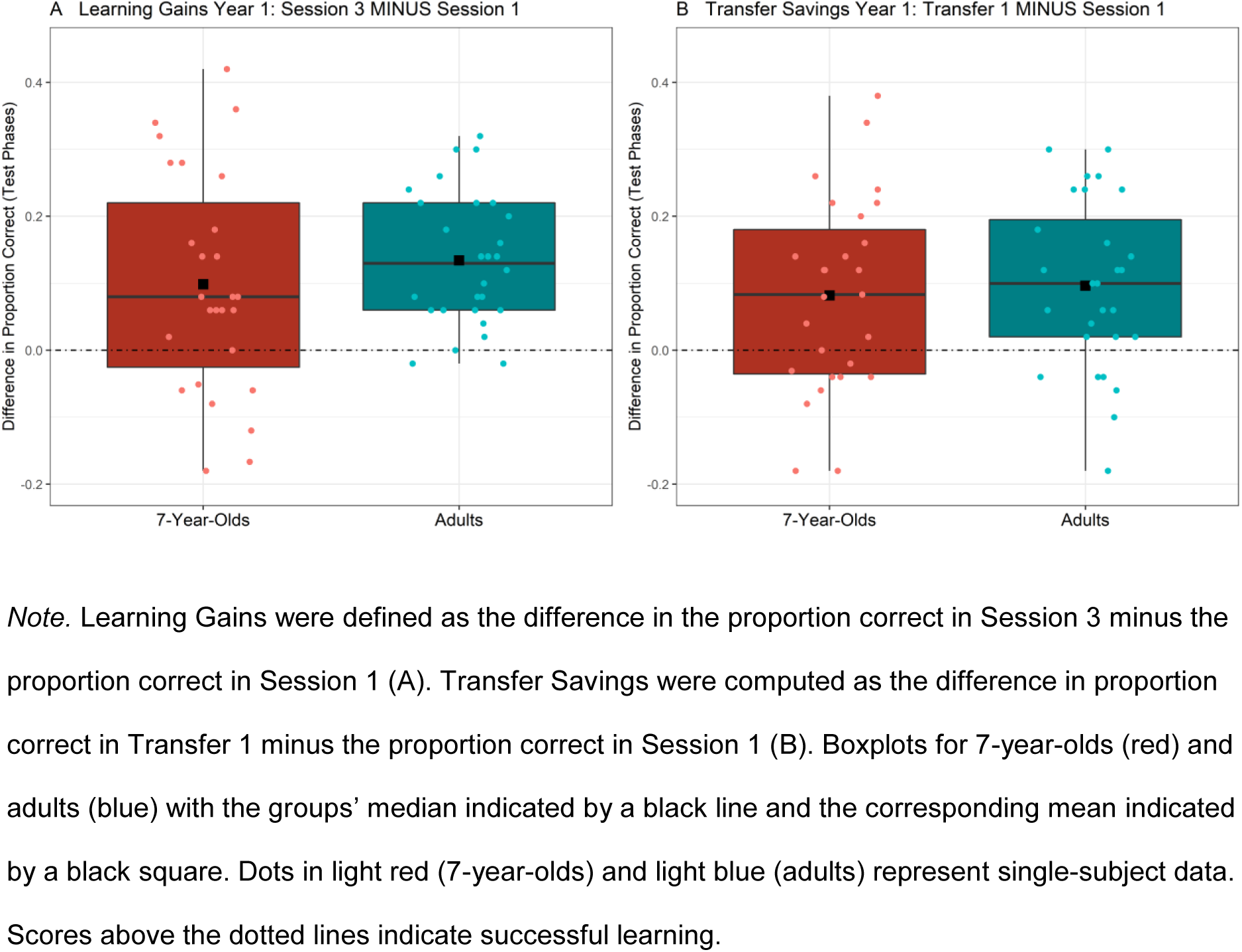
Learning Gains and Transfer Savings in Year 1.

Next, we asked whether children and adults were able to apply their acquired rule knowledge of the first three sessions to the second stimulus set in the transfer session (Transfer 1; see Fig. 4B). To examine transfer savings to the new stimulus set, mean session scores were entered into an ANOVA with the factors *Age* (between-subject; levels: 7-year-olds, adults) and *Session* (within-subject; levels: Session 1, Transfer 1). Adults outperformed children (see Table 2), irrespective of the session (main effect of *Age*: *F*(1, 53) = 180.35, *p* < .001, η^2^_g_ = .64; *BF_incl_* > 100). Crucially, in both groups performance in the transfer session exceeded performance in the first session (main effect of *Session: F*(1, 53) = 23.70, *p* < .001, η^2^_g_ = .18; *BF_incl_* > 100), indicating successful transfer to the new stimulus set (see Fig. 4B). This transfer effect did not significantly differ in size between adults and 7- year-olds (interaction *Age*Session*: *F*(1, 53) = 0.16, *p* = .694, η^2^_g_ < .01; *BF_incl_* = 0.35).

We further tested whether performance changed from the last session with the first stimulus set (Session 3) to the subsequent transfer session (Transfer 1): An ANOVA with the factors *Age* (between-subject; levels: 7-year-olds, adults) and *Session* (within-subject; levels: Session 3, Transfer 1) revealed a significant main effect of *Age* (*F*(1, 53) = 106.37, *p* < .001, η^2^_g_ = .60; *BF_incl_* > 100) with adults performing better than 7-year-olds across all sessions (see Table 2). No significant performance loss (main effect of *Session*: *F*(1, 53) = 3.08, *p* = .085, η^2^_g_ = .01; *BF_incl_* = .74) or age difference in their preserved performance level in the transfer session (interaction *Age*Session F*(1, 53) = 0.47, *p* = .498, η^2^_g_ < .01; *BF_incl_* = .31) was observed.

Taken together, across one week in Year 1, both children and adults learned the AG and finally successfully applied this knowledge to a new stimulus set. Throughout the sessions, adults performed at a higher level than children while across-session learning gains did not differ between groups.

### Identifying the First Learning Trial from Modeling trial-by-trial Performance

For within-session learning, the state-space model by Smith et al. (2005) identified the very first test trial (i.e., in the beginning of Block 1 in Session 1) in the adult group as the first timepoint at which learning had taken place. This means that adults showed within-session learning effects after being exposed to a single learning phase of 18 grammatical sequences. In contrast, for children, the 30^th^ test trial (i.e., at the end of Block 3 in the second half of Session 1) was identified as the first timepoint when learning had happened. Thus, children needed more learning trials until they featured successful learning of the AG in Year 1.

### Performance Correlations with Explicit Sequence Knowledge and Cognitive Skills

When comparing the two age groups, scores of reported explicit sequence knowledge did not significantly differ between 7-year-olds and adults (*t*(53) = -1.17, *p* = .249, *d* = 0.31, *BF_10_* = .48). In 7-year-olds, explicit knowledge about sequence rules (assessed at the end of Transfer 1) was marginally positively correlated with Transfer Savings (Transfer 1 – Session 1) (*r_s_* = .42, *p* = .058, *BF_10_* = 1.68; adults: *r_s_* = .25; *p* = .193, *BF_10_* = .55). Learning Gains (Session 3 – Session 1) were positively associated with working memory capacity in 7-year-olds (*r_s_* = .57, *p* = .004, *BF_10_* = 24.46), but not in adults (*r_s_* = .04, *p* = .849, *BF_10_* = .25). As seen from Appendix D (Table D.1) other associations between explicit sequence knowledge, memory and language skills and AGL performance were not statistically significant (all | *r_s_* | ≤ .32, all *p* ≥ .172, all *BF_10_* ≤ 1.11).

### Relearning after a One-Year-Delay (Year 1 vs. Year 2)

The following analyses were performed for the subgroup of 16 seven-year-olds and 20 adults, who completed the four home follow-up sessions in Year 2 in addition to all sessions in Year 1. Thus, we first tested, whether this subgroup performed above chance in both Year 1 and Year 2 (see black group means in Fig. 5). All following analyses are based group *Means(SDs)* in Table 2 (see data for *n* = 16 seven-year-olds & *n* = 20 adults).

**Figure 5.**
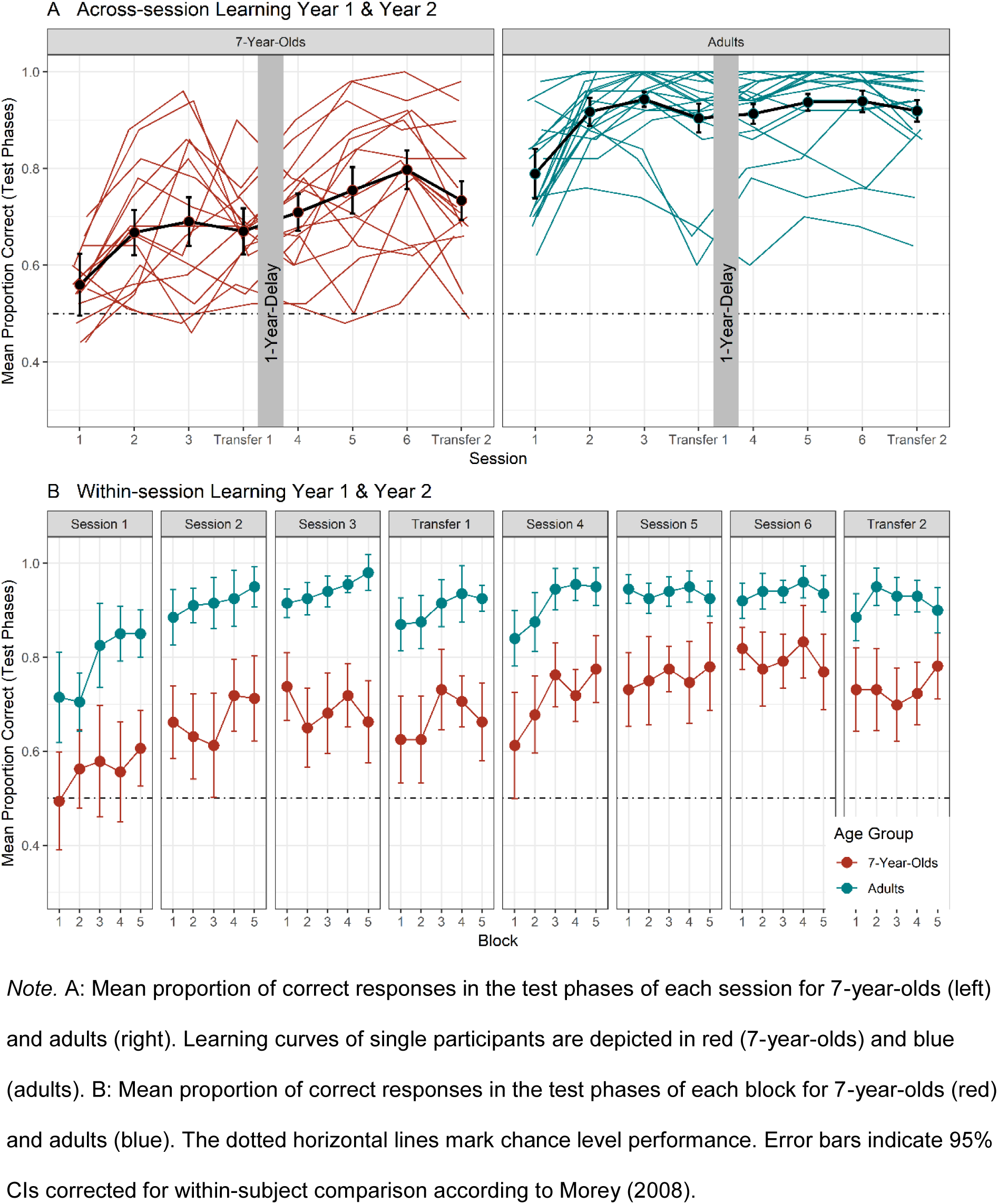
Performance Trajectories Across Sessions and Within Sessions of Year 1 & Year 2.

For all eight sessions, group-level performance exceeded the chance level of 0.5 (two-alternative forced-choice trials) in 7-year-olds (all *t*(15) ≥ 3.30, all *p* ≤ .008, all *d* ≥ 0.83, all *BF_+0_* > 100; one-sided) and in adults (one-sided; all *V*(19) = 210.00, all *p* ≤ .008, all *r* = 0.88, all *BF_+0_* > 100; one-sided).

### Initial and Final Performance Levels in Year 1 vs. Year 2

To evaluate gains for relearning in Year 2 from the previous learning experience in Year 1, we tested mean performance in the very first session (Session 1) against mean performance in the first session of relearning in Year 2 (Session 4) (see Fig. 6A): An ANOVA with the factors *Age* (between-subject; levels: 7-year-olds, adults) and *Session* (within-subject; levels: Session 1, Session 4) revealed higher overall performance levels in adults than in children (see Table 2; main effect of *Age F*(1, 34) = 88.02, *p* < .001, η^2^_g_ = .57; *BF_incl_* > 100). Importantly, both age groups performed better in the first relearning session of Year 2 than in the first session of Year 1 (see Table 2; main effect of *Session F*(1, 34) = 36.92, *p* < .001, η^2^_g_ = .35; *BF_incl_* > 100), as indicated by the positive mean difference scores (see Figure 7A). The performance gain at relearning in Year 2 was of similar magnitude for 7-year-olds and adults (*Age*Session* interaction *F*(1, 34) = 0.33, *p* = .569, η^2^_g_ < .001; *BF_incl_* = .35).

**Figure 6.**
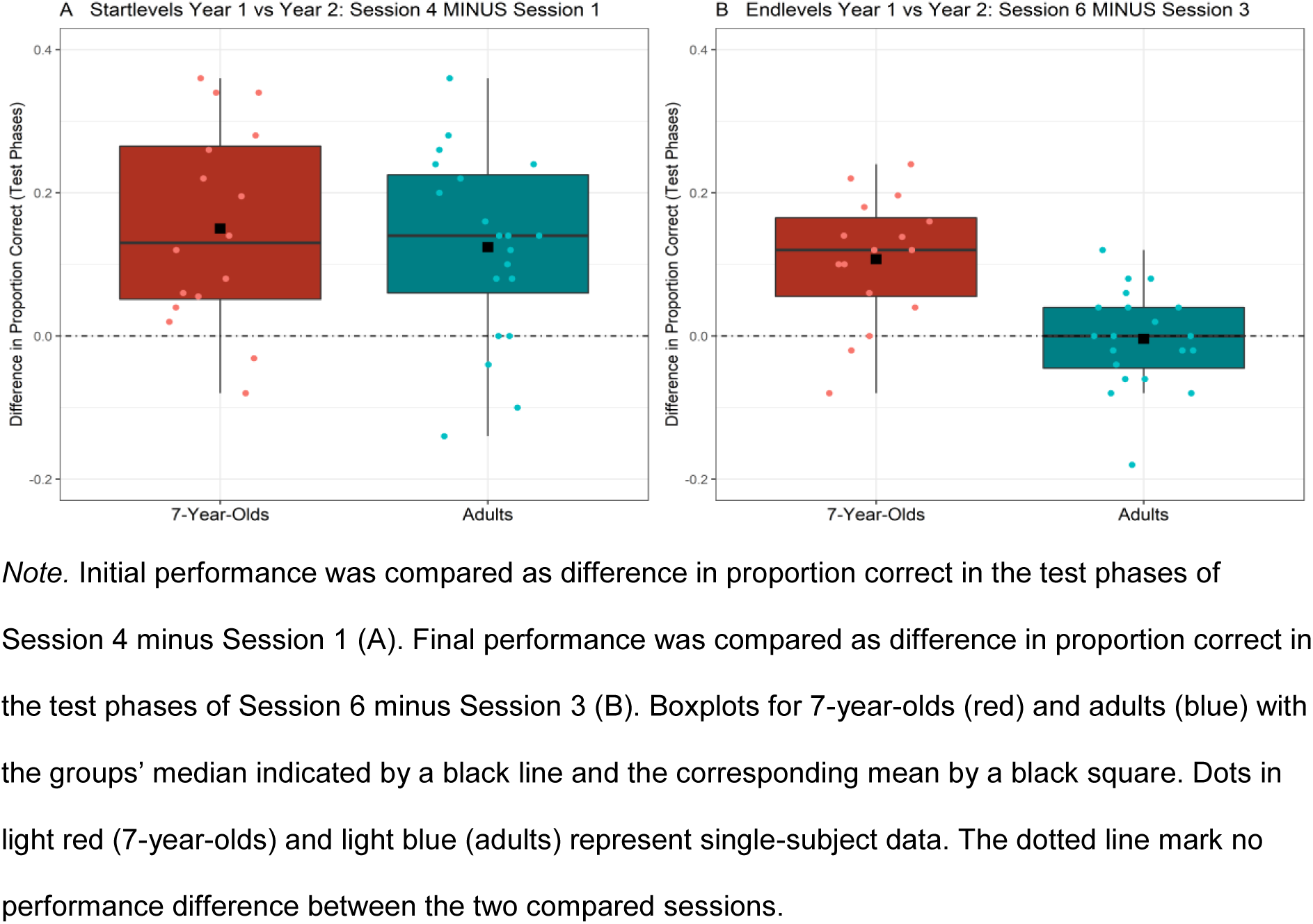
Session Differences of Year 1 & Year 2 for Initial and Final Performance Levels.

**Figure 7.**
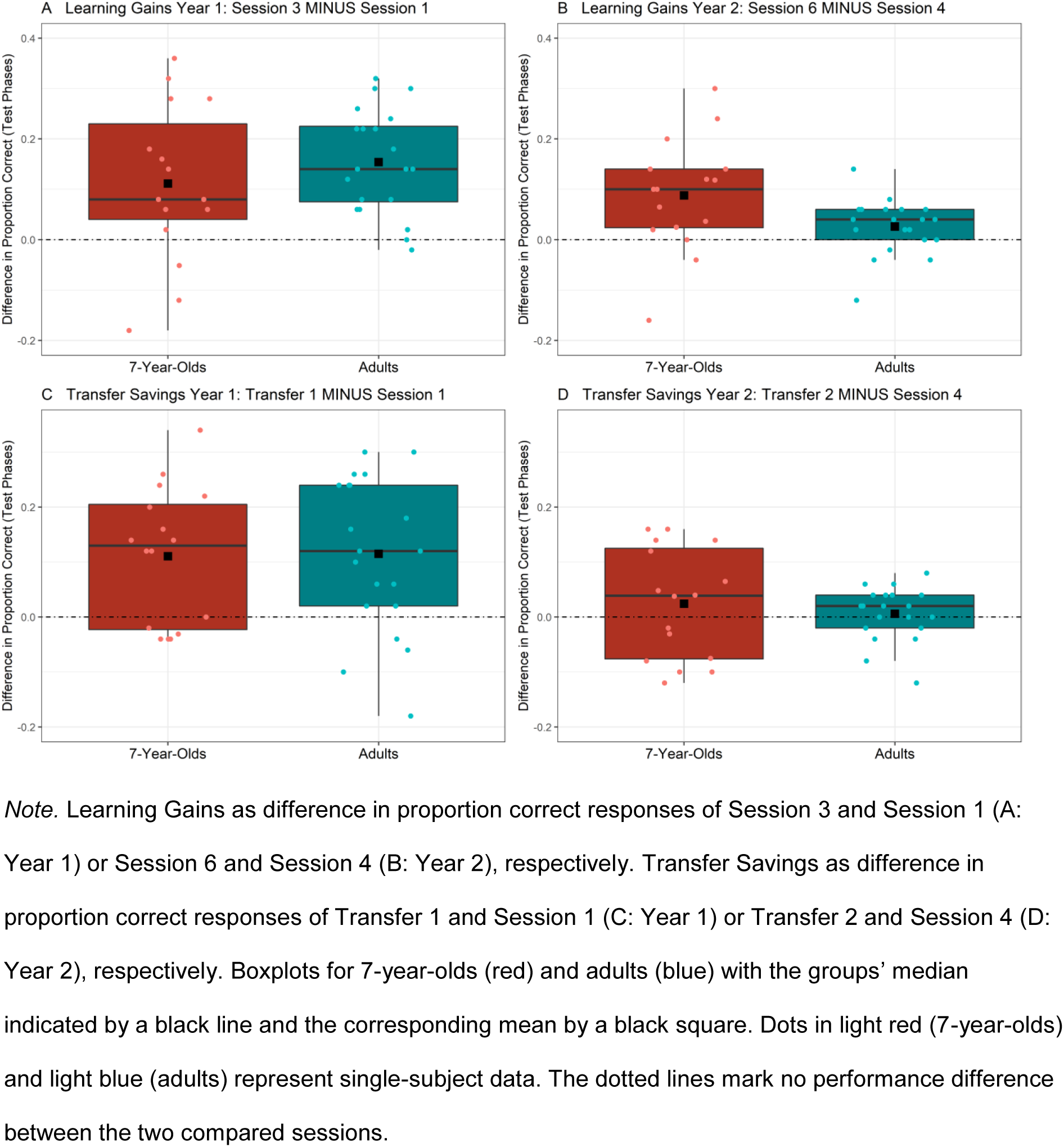
Session Differences for Learning Gains and Transfer Savings for Year 1 & Year 2.

Next, we tested whether both age groups had lost in performance from the last session with the first stimulus set (Session 3) in Year 1 to the first session of relearning of Year 2 (Session 4). An ANOVA with the factors *Age* (between-subject; levels: 7-year-olds, adults) and *Session* (within-subject; levels: Session 3, Session 4) revealed only a significant main effect of *Age* (*F*(1, 34) = 42.81, *p* < .001, η^2^_g_ = .52; *BF_incl_* > 100), with adults performing better than 7-year-olds, irrespective of the session (see Table 2). Participants retained their performance level from the end of Year 1 in Year 2 (main effect of *Session*: *F*(1, 34) = 0.11, *p* = .740, η^2^_g_ < .01; *BF_incl_* = .25), with no significant age difference in adults’ and children’s retention (interaction *Age*Session F*(1, 34) = 2.43, *p* = .128, η^2^_g_ = .01, *BF_incl_* = 0.81).

We next examined whether relearning in Year 2 resulted in a higher final performance level than in Year 1. Mean performance in the last session with the first stimulus material was compared between Session 3 and Session 6 (see difference scores for both age groups see Fig. 6B). An ANOVA with the factors *Age* (between-subject; levels: 7-year-olds, adults) and *Session* (within-subject; levels: Session 3, Session 6) yielded significant main effects of *Age* (*F*(1, 34) = 28.17, *p* < .001, η^2^_g_ = .42; *BF_incl_* > 100) and *Session* (*F*(1, 34) = 15.37, *p* < .001, η^2^_g_ = .05; *BF_incl_* = 5.29), as well as a significant interaction of *Age*Session* (*F*(1, 34) = 17.85, *p* < .001, η^2^_g_ = .06; *BF_incl_* = 83.03): Only 7-year-olds performed better in Session 6 of Year 2, compared to Session 3 of Year 1 (see Table 2; *t*(15) = -4.83, *p* < .002, *d* = 1.21, *BF_10_* > 100; shown in red in Fig. 6B left panel), while adults’ final performance did not differ for Session 3 of Year 1 and Session 6 of Year 2 (see Table 2; *t*(19) = 0.26, *p* = .799, *d* = 0.06, *BF_10_* = .24; shown in blue in Fig. 6B right panel).

### Performance Improvement and Transfer in Year 1 vs. Year 2

To compare performance improvements over three sessions with the first stimulus set in both years, we tested initial learning in Year 1 (see Fig. 7A) against relearning in Year 2 (see Fig. 7B): An ANOVA with the factors *Age* (between-subject; levels: 7-year-olds, adults),*Year* (within-subject; levels: Year 1, Year 2) and *Session* (within-subject; levels: First Session [Year 1: Session 1; Year 2: Session 4], Last Session[Year 1: Session 3; Year 2: Session 6]) revealed significant two-way interactions of *Age*Year* (*F*(1, 34) = 7.83, *p* = .008, η^2^_g_ = .03; *BF_incl_* = 3.26) and *Year*Session* (*F*(1, 34) = 12.79, *p* = .001, η^2^_g_ = .04; *BF_incl_* = 23.13) as well as main effects of *Age*, *Year* and *Session* (all *F*(1, 34) ≥ 25.46, all *p* < .001, all η^2^_g_ ≥ .11, all *BF_incl_* > 100; all other *F*(1, 34) ≤ 3.16, *p* ≥ .085, η^2^_g_ ≤ .01, *BF_incl_* < .69): Overall, participants improved to a greater degree over three sessions in Year 1 (Session 1 vs. Session 3, see Fig. 7A) than they did in Year 2 (Session 4 vs. Session 6, see Fig. 7B; performance improvement in Year 2 [Session 6 – Session 4] vs. performance improvement in Year 1 [Sessions 3 - 1]: *t*(35) = 3.63, *p* < .002, *d* = 0.60, *BF_10_* = 34.34), and 7-year-olds in general gained more from learning in Year 2 than adults (performance difference average Year 2 [Sessions 4 & 6] – average Year 1 [Sessions 1 & 3] in children vs. adults: *t*(34) = 2.47, *p* = .019, *d* = 0.83, *BF_10_* = 3.12).

To compare how participants transferred learned regularities to a second stimulus set in Year 1 versus Year 2 (see Fig. 7C & 8D), we conducted an ANOVA with the factors *Age* (between-subject; levels: 7-year-olds, adults),*Year* (within-subject; levels: Year 1, Year 2) and *Session* (within-subject; levels: First Session [Year 1: Session 1; Year 2: Session 4], Transfer Session [Year 1: Transfer 1; Year 2: Transfer 2]). This analysis revealed a significant interaction of *Year*Session* (*F*(1, 34) = 13.83, *p* = .001, η^2^_g_ = .06*; BF_incl_* = 73.70 and significant main effects of *Age, Year* and *Session* (all *F*(1, 34) ≥ 25.30, all *p* < .001, all η^2^_g_ ≥ .09, all *BF_incl_* > 100; all other *F*(1, 34) ≤ 1.61, *p* ≥ .214, η^2^_g_ ≤ .01, *BF_incl_* ≤ .52): Overall, performance in Transfer 1 exceeded performance in Session 1 in Year 1 (*t*(35) = -5.14, *p* < .002, *d* = 0.86, *BF_10_* > 100; see Fig. 7C), but this was not the case in Year 2 for Transfer 2 compared to Session 4 (*t*(35) = -1.11, *p* = .274, *d* = 0.19, *BF_10_* = .32; see Fig. 7D). This effect was similar in size for both age groups (*Age*Year*Session: F*(1, 34) = 0.18, *p* = .671, η^2^_g_ < .01; *BF_incl_* = .34), Thus, no additional performance benefit for the second stimulus set was observed after relearning in Year 2.

We further investigated preserved performance in the transfer session compared to the directly preceding session with the first stimulus set in Year 1 versus Year 2. To this end, we conducted an ANOVA with the factors *Age* (between-subject; levels: 7-year-olds, adults), *Year* (within-subject; levels: Year 1, Year 2) and *Session* (within-subject; levels: Last Session [Year 1: Session 3; Year 2: Session 6], Transfer Session [Year 1: Transfer 1; Year 2: Transfer 2]). This analysis yielded a significant two-way interaction of *Age*Year* (*F*(1, 34) = 16.86, *p* < .001, η^2^_g_ = .03; *BF_incl_* = 43.97) in addition to significant main effects of *Age, Year* and *Session* (all *F*(1, 34) ≥ 8.26, all *p* ≤ .007, all η^2^_g_ ≥ .03, all *BF_incl_* ≥ 14.18; all other *F*(1, 34) ≤ 1.95, *p* ≥ .171, η^2^_g_ < .01, *BF_incl_* ≤ .70): 7-year-olds displayed performance gains from Year 1 to Year 2 (Year 1 vs. Year 2: *t*(31) = -4.51, *p* < .002, *d* = 0.80, *BF_10_* > 100), while adults did not (Year 1 vs. Year 2: *t*(39) = -0.49, *p* = .623, *d* = 0.08, *BF_10_* = .25). For both age groups, performance declined from the last session (i.e., average of Session 3 & 6) to the transfer session (i.e., average of Transfer 1 & 2; main effect of *Session: F*(1, 34) = 8.26, *p* = .007, η^2^_g_ = .03, *BF_incl_* = 14.18). No difference in this second transfer contrast emerged when comparing Year 1 and Year 2 (*Year*Session: F*(1, 34) = 0.30, *p* = .596, η^2^_g_ < .01; *BF_incl_* = .25), independently of age (*Age*Year*Session: F*(1, 34) = 1.95, *p* = .171, η^2^_g_ < .01; *BF_incl_* = .70). These non-significant interactions together with the significant main effect of *Session* indicate that the performance of participants of both age groups decreased from the last session with the first stimulus set to the transfer session in both Year 1 and Year 2.

In summary, both age groups preserved their performance from Session 3 (last session) of Year 1 to Session 4 (first session) of Year 2 and both groups performed higher in the first session of Year 2 compared to the first session of Year 1. Both groups improved more over the three sessions (with stimulus-set-1) in Year 1 than in Year 2, and showed no additional transfer benefits for the second stimulus set in Year 2. Seven-year-olds, but not adults, reached higher final performance levels with the first stimulus set after relearning in Year 2, compared to Session 3 (last session) in Year 1. Adults performed close to ceiling in Year 1 already.

### Identifying the First Learning Trial from Modeling trial-by-trial Performance

For within-session learning in Year 2, the state-space model by Smith et al. (2005) identified the very first test trial (Block 1 in Session 1) in both age groups as the timepoint at which learning first happened. This means that at relearning, both groups showed within-session learning effects after being exposed to a single learning phase of 18 grammatical sequences. Thus, children demonstrated within-session learning effects at relearning in Year 2 as early as adults.

### Performance Correlations with Explicit Sequence Knowledge and Cognitive Skills

Scores in reported explicit sequence knowledge in Year 2 (assessed in the end of Transfer 2) did not significantly differ between 7-year-olds and adults (*t*(34) = 0.43, *p* = .249, *d* = 0.18, *BF_10_* = .35). 7-year-olds reported more explicit knowledge about sequence rules at the end of Year 2, compared to the end of Year 1 (*t*(15) = -3.51, *p* = .004, *d* = 0.90, *BF_-0_* = 56.33; one-sided), while this difference between Year 1 and 2 did not reach statistical significance in adults (*t*(19) = -1.43, *p* = .084, *d* = 0.32, *BF_-0_* = 1.02; one-sided). As seen from Appendix D (Table D.2) none of the associations between explicit sequence knowledge (Year 1 & Year 2) or memory and language skills (Year 1) with AGL performance (Year 2), respectively, reached statistical significance (all *r_s_* ≤ .49, all *p* ≤ .114, all *BF_10_* ≤ 2.45).

## Discussion

The goal of the present study was to test whether 7-year-olds outperform adults in artificial grammar learning and whether this advantage translates to superior transfer and long-term memory of AG rules. To this end, both 7-year-olds and adults engaged in a multisession visual AGL task which was repeated one year later.

We found successful AG learning and transfer of rule knowledge to another stimulus set in both 7-year-olds and adults. However, adults learned quicker and overall performed at a higher level. Both groups retained rule knowledge over a one-year-period, started at the level reached one year earlier and continued to improve, although to a lesser degree than in the first year. We did not find better retention or larger relearning advantages in 7-year-olds compared to adults in Year 2. Explicit knowledge of the AG rules was indistinguishable between adults and children and correlated with transfer gains in children.

### 7-year-olds acquire AG rules but overall learn slower and perform worse than adults across several sessions

When exposed to a visual AG, 7-year-olds and adults demonstrated successful learning of the AG. Adults overall outperformed children, but learning gains over several sessions were indistinguishable between both groups. These results are in accord with Ferman and Karni (2010) and Smalle, Page, et al. (2017), who reported indistinguishable learning rates across multiple sessions in 8-12-year-olds and adults in phonological sequence learning tasks. Our study extends these findings to visual sequence learning with more complex rules as defined in an AG.

Our study allowed us to directly compare both age groups in short-term retention, which elaborates evidence on visuomotor retention across 24 hours in an alternating serial reaction time task for 9-15-year-old children (Tóth-Fáber et al., 2021) and adults (Kóbor et al., 2017) from separate studies. Our finding that 7-year-olds and adults continued learning at their last performance level after delays of at least one night between the sessions of Year 1 suggests similarly effective short-term consolidation processes for learned regularities in children and adults. This is in line with a recent study (Tóth-Fáber et al., 2023) in which the same authors followed up on their previous findings (Kóbor et al., 2017; Tóth-Fáber et al., 2021); in a large sample of age 7- to 76-year-old participants, they did not find any age-related differences in the 24-hour-consolidation of visuomotor learning.

In contrast to previous findings on rule generalization in phonological single-session (Hickey et al., 2019) and multi-session (Ferman & Karni, 2010) learning, we observed that children as young as seven years generalized their visual rule knowledge to the same extent as adults did. Transfer effects in our study emerged without any explicit instruction, that is, without providing participants explanations about the nature of these rules. Our finding, thus, is incompatible with the proposition that explicit instructions are necessary for successful transfer in children younger than 12 years (Ferman & Karni, 2010, 2014). In line with our results, in a single-session study, 6- to 9-year-olds implicitly extracted categorical regularities in visual triplet learning and transferred their rule knowledge to unseen items of the same picture category to the same degree as adults (Jung et al., 2020). Even younger children with age 3 to 6 years were reported to successfully transfer learned regularities to new syllables in auditory artificial grammar learning if underlying sequence rules resembled such occurring in natural language (Nowak & Baggio, 2017). These inconsistent findings raise the question of under which conditions children transfer learned regularities to new items or even to new categories. In line with previous suggestions (Gomez, 1997; Nowak & Baggio, 2017; Witt et al., 2013), we hypothesize that in learning complex regularities, task features that help building explicit knowledge about these regularities, such as providing performance feedback at test (Nowak & Baggio, 2017), might favor rule abstraction and consequently rule transfer to new surface features in children. Becoming aware of underlying sequence rules, in turn, has been proposed to drive the consolidation of rule knowledge in an offline period with sleep (Janacsek & Nemeth, 2012).

A link between explicit rule knowledge and rule transfer is supported by our finding that in children larger Transfer Savings (Transfer 1 minus Session 1) showed a trend to be positively associated with more explicit knowledge of the rules at the end of Year 1. Furthermore, children benefitted from higher working memory capacities for their Learning Gains in the first Year (Session 1 to 3) suggesting that the ability to explicitly represent sequences in short-term memory contributed to their learning in the current task. In fact, short term memory has been discussed as a gating factor in learning sequential regularities, especially for non-adjacent regularities (Conway, 2020).

Despite their similarities in learning trajectories and transfer effects, children and adults displayed differences across the three sessions of Year 1. Adults needed less exposure to the AG within the first session for initial learning: After only one learning phase with 18 grammatical sequences they performed above-chance during test. By contrast, successful learning in children was not observed before the third test phase of Session 1, i.e., only after exposure to 54 grammatical sequences. The statistical learning model of Janacsek et al. (2012) (further elaborated by Daltrozzo & Conway, 2014; Janacsek et al., 2012; Nemeth et al., 2013) proposes that in the course of development, a switch to a more supervised, “model-based” learning system takes place. According to these authors, model-based learning relies more on attentional resources, behavioral control and prior knowledge than the “model-free” learning It might be speculated that the stronger reliance on model-based learning might have contributed to the superior performance of adults compared to 7-year-olds in the present task setting. However, children and adults reported indistinguishable levels of rule knowledge in Year 1 and Year 2. This finding implies that at least the accessible rule representations as a result of learning were similar in both age groups. Future neuroimaging studies would allow to more directly identify the age-dependent mechanisms of repeated learning.

The lack of a significant association between explicit knowledge and sequence learning in adults might be due to ceiling performance in this group. In fact, ceiling performance in adults was observed as early as Session 2. However, adults and children in the present study did not significantly differ in their reported levels of explicit sequence knowledge by the end of the transfer session. This might be due our late assessment of explicit sequence knowledge after four learning sessions, in order to avoid inducing a change in learning strategies. Wilhelm et al. (2013) tested the explicit recall of adjacent items from learning a deterministic sequence in an implicit visuomotor task after a sleep vs. wake phase. They found that children aged 8-11 years benefitted more than adults from sleep in acquiring explicit knowledge from their implicit learning experience. Based on this finding, it can be hypothesized that our study design aided the formation of explicit knowledge about sequence rules especially in children, possibly mitigating previously reported age differences in explicit rule knowledge.

### Children and adults retain visual regularities over a one-year delay and show comparable relearning effects after the delay

Both age groups by and large retained rule knowledge over the 12-month break. This corroborates findings from phonological rule learning which suggested long-term retention of sequence knowledge across two months (Ferman & Karni, 2010) and one year (Smalle, Page, et al., 2017). These studies included both 8- to 12-year-old children and adults. Our results on retaining complex visual regularities furthermore extend findings on visuomotor retention across a one year delay on 9-15-year-old’s (Tóth-Fáber et al., 2021) and adults’ learning (Kóbor et al., 2017) from separate studies: We were able to directly compare children and adults and show that adult-like retention is already evident in children as young as 7 years.

In addition to testing retention of complex visual regularities after a long-term delay in a single follow-up session, we additionally identified how both age groups use their acquired rule knowledge in another set of multiple relearning sessions in Year 2: Children and adults continued to improve with the first stimulus set, although to a lesser degree than in Year 1. This was true for both age groups but more so for children, because adults performed close to ceiling from Session 2 in Year 1 onwards.

We cannot exclude the possibility that in Year 2, children’s performance was influenced by age-related better sequence learning abilities compared to Year 1, e.g., due to a higher memory capacity. Previous studies (Arciuli & Simpson, 2011; Raviv & Arnon, 2017; Shufaniya & Arnon, 2018) have demonstrated that children’s behavioral performance in visual triplet learning tasks improves between 5 and 12 years of age. Available data for 6.5-7.5-year-olds vs. 8-9-year-olds from the cross-sectional study by Raviv and Arnon (2017) showed inconsistent trends for different stimulus materials, with younger children performing better in auditory/linguistic triplet learning vs. older children performing better in visual/non-linguistic triplet learning. We think that two observations speak against the notion that relearning effects in children reported here are due to maturation: First, children preserved their last performance level across one year (Session 3 to Session 4), corroborating the phenomenon of experience-dependent benefits for relearning in school-aged children reported in previous studies (Smalle, Page, et al., 2017; Tóth-Fáber et al., 2021). Second, the relearning gain with the same stimulus material over three sessions in Year 2 (Session 4 to 6) compared to the learning rate in Year 1 (Session 1 to 3) was lower. If children were better at acquiring regularities due to their more matured cognitive abilities in Year 2, they would have been expected to show a greater increase in learning in Year 2 compared to Year 1. Importantly, all relearning effects were confirmed after eliminating the first task block of Session 1 of Year 1 and Session 4 (first session) of Year 2, respectively, to exclude trivial task familiarity effects (Appendix C, Table C.1-3).

Both age groups did not show additional improvements in the transfer to the second stimulus set in Year 2. Since additional experience with the first stimulus material further improved performance in Year 2 (Session 4 to 6), it seems unlikely that additional transfer gains in Year 2 (Session 4 to Transfer 2) were prevented by ceiling effects. In general, offline periods between learning sessions seem to promote the extraction and representation of underlying regularities by replay-induced strengthening of memory representations at the level of neural circuits (Lerner & Gluck, 2019; Liu et al., 2019; Wilhelm et al., 2012). It could be speculated that replay or similar mechanisms enabled generalized representations to be formed from spaced learning in Year 1, already. Further research should clarify neural mechanisms of repeated learning across extended time periods like months, e.g. with regard to structural adaptations shown in non-human animal models of complex skill learning (Hofer & Bonhoeffer, 2010).

## Summary and conclusion

In summary, the present study demonstrated successful artificial grammar (AG) learning and transfer of rule knowledge to new visual stimuli in both 7-year-olds and adults; Adults learned quicker and overall performed higher. Both age groups successfully applied retained rule knowledge after one year without any advantage for either group. Explicit knowledge of the AG was indistinguishable between adults and children and correlated with transfer performance in children. These findings challenge the hypothesis that learning advantages in mid childhood result from higher implicit rule learning and retention capacities.

## Acknowledgements

We thank Leon Bauer, Maike Fuchs, Nicola Kaczmarek, Kathrin Lambeck, Alina Scheller, Dagmar Tödter, and Jelka Wöbke and for help in participant recruitment and data collection. We are grateful to Veronika Zweckerl for comments on the study design and Liesa Stange and Alexander Kramer for their help in setting up the data analysis pipeline. Special thanks goes to the families for participating in a time consuming study.

## Author Contributions

Daniela Schönberger: Conceptualization, Methodology, Data curation, Software, Formal analysis, Visualization, Writing – original draft, Project administration; Patrick Bruns: Conceptualization, Methodology, Supervision, Writing – review and editing, Brigitte Röder: Conceptualization, Methodology, Resources, Supervision, Funding acquisition, Writing – review and editing.

