## Supplemental Material for "Visual Artificial Grammar Learning Across One Year in 7-Year-Olds and Adults"

### Appendix A.

#### Participant Characteristics

**Table A.1**

*Participant Characteristics of the Final Sample for Analyses of Year 1 across 1 Week*

| **Participant Characteristics** | **7-Year-Olds (*n* = 27)** | **Adults (*n* = 28)** |
|---|---|---|
| Days between Sessions | | |
| Session 1 to 3 | 4.11 (1.19) | 4.00 (1.59) |
| Session 1 to Transfer 1 | 7.00 (0.73) | 6.75 (1.08) |
| Age (years) | 7.05 (0.07) | 23.12 (3.50) |
| Age span | 6.91 – 7.20 | 18.83 – 33.54 |
| Gender (f/m) | 15/12 | 18/10 |
| School/Education [a] | *n* = 26 1st grade elementary school<br>*n* = 2 2nd grade elementary school | *n* = 28 university students |
| Bilinguals | 4 | 6 * |
| Daily mobile device usage (min) [a] | 9.07 (15.45) | 254.77 (126.00) |
| Explicit knowledge Year 1 [b] | 0.46 (0.23) | 0.54 (0.26) |

*Note. M (SD).*
[*] *n* = 25 (3 missing values).

[a] assessed in Year 1 (Session 1).

[b] scores could range from -1 (no rule knowledge) to 1 (max. rule knowledge).

**Table A.2**

*Participant Characteristics of the Final Sample for joint Analyses of Year 1 & Year 2*

| **Participant Characteristics** | **7-Year-Olds (*n* = 16)** | **Adults (*n* = 20)** |
|---|---|---|
| Time period betw. Year 1 & 2 (months betw. Session 1 & 4) | 13.00 (1.10)<br>[11.00-15.00] | 12.75 (0.44)<br>[12.00-13.00] |
| Days between Sessions of Year 1 | | |
| Session 1 to 3 | 4.19 (0.83) | 3.80 (1.54) |
| Session 1 to Transfer 1 | 6.88 (0.50) | 6.60 (1.14) |
| Days between Sessions of Year 2 | | |
| Session 4 to 6 | 2.63 (0.81) | 2.45 (0.60) |
| Session 4 to Transfer 2 | 4.19 (1.52) | 3.85 (0.93) |
| Age Year 1 (years) | 7.06 (0.08)<br>[6.92-7.20] | 22.14 (2.48)<br>[18.83 – 27.36] |
| Age Year 2 (years) | 8.18 (0.09)<br>[8.03-8.31] | 23.24 (2.48)<br>[19.94 – 28.47] |
| Gender (f/m) | 12/4 | 14/6 |
| Education [a] | *n* = 15 1st grade elementary school<br>*n* = 1 2nd grade elementary school | *n* = 20 university students |
| Bilinguals | 3 | 6 * |
| Daily mobile device usage (min) [a] | 9.24 (15.46) | 250.96 (120.89) |
| Explicit knowledge Year 1 [b] | 0.51 (0.17) | 0.57 (0.26) |
| Explicit knowledge Year 2 [b] | 0.68 (0.15) | 0.66 (0.21) |

*Note. M (SD)* [range], betw. = between.
[*] *n* = 17 (3 missing values).

[a] assessed in Year 1 (Session 1).

[b] scores could range from -1 (no rule knowledge) to 1 (max. rule knowledge).

**Appendix B.**

**Stimulus Materials**

**Table B.1**

*Picture Assignment in AGL Task, corresponding to Numbers in Figure 2*

| # in Fig. 2 | Assigned picture Stimulus Set Animals, Version 1 | Assigned picture Stimulus Set Animals, Version 2 | Assigned picture Stimulus Set Colors, Version 1 | Assigned picture Stimulus Set Colors, Version 2 |
|---|---|---|---|---|
| 1 | 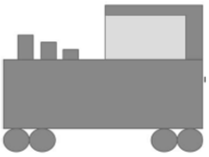 | 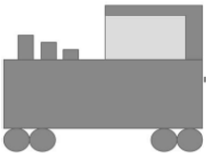 |  |  |
| 2 | 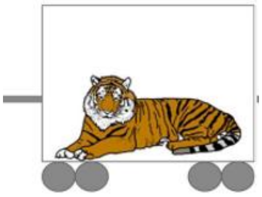 | 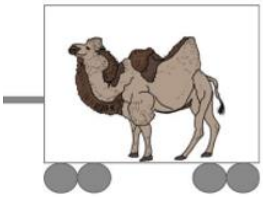 | 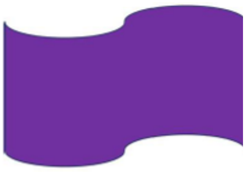 | 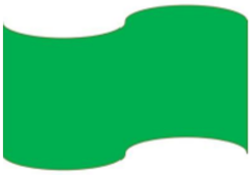 |
| 3 | 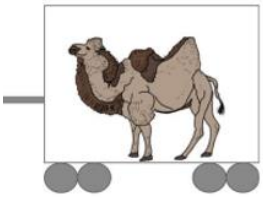 | 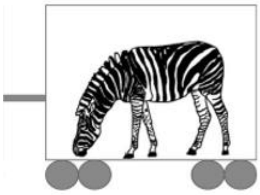 | 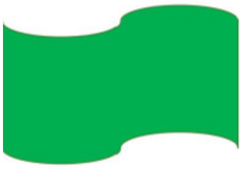 | 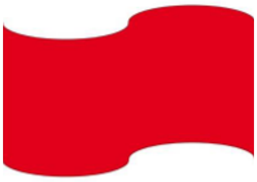 |
| 4 | 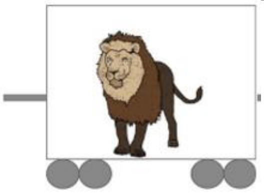 | 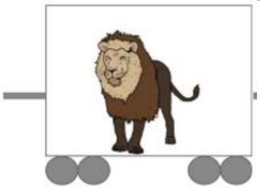 | 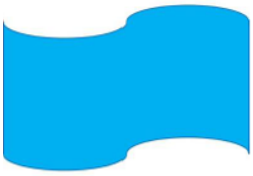 |  |
| 5 | 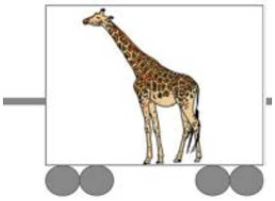 | 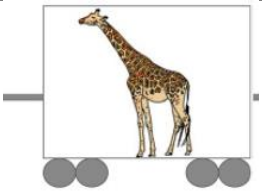 |  | 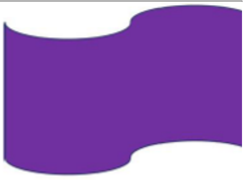 |
| 6 | 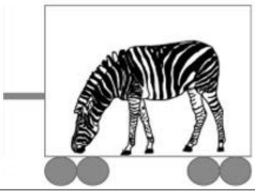 | 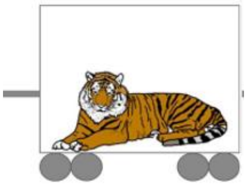 | 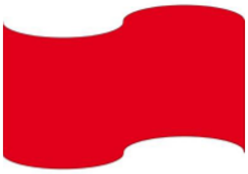 | 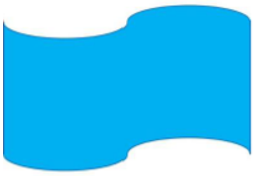 |

*Note.* Picture assignments for both task versions per stimulus set were randomly picked from all possible assignments from permutations generated by a randomization script. The resulting task versions were applied in a counterbalanced manner in both age groups.

**Table B.2**

*Instructions for the AGL task*

| **Stimulus Set** | **Learning Phase** | **Test Phase** | **Each Test Trial** |
|---|---|---|---|
| Animals | ‘‘Shortly, you will see Ms. Pepe´s circus. She travels along the country with her circus in a train. There is one animal in each car of the train. Ms. Pepe arranges the animals in such a way that they feel comfortable with each other. You are a detective who will see several of Ms. Pepe´s trains. Watch them carefully, we will ask questions about them afterwards.” | ‘‘You are a detective who arrives at the station and finds two circus trains there. Only one of them belongs to Ms. Pepe. Help us identify which one belongs to her. Use your gut feeling and choose the train that first comes to your mind.<br>To choose a train, just tap it on the screen.” | ‘‘Which circus train was arranged by Ms. Pepe? Tap it on the screen.” |
| Colors | ‘‘Shortly, you will see flags of the sports team *Strong Tigers*. The flags are made from different colors and are used at sports tournaments. To each tournament, the *Strong Tigers* bring a different flag, so their fans won’t be bored. They arrange their flags in a way so the colors go along well with each other. You are a detective who will see several of the *Strong Tiger’s* flags. Watch them carefully, we will ask questions about them afterwards.” | ‘‘You are a detective who arrives at a tournament and finds two team flags there. Only one of them belongs to the *Strong Tigers*. Help us identify which one belongs to them. Use your gut feeling and choose the flag that first comes to your mind.<br>To choose a flag, just tap it on the screen.” | ‘‘Which flag was arranged by the *Strong Tigers*? Tap it on the screen.” |

*Note*. Translated from the original instructions which were in German. A shortened version of the learning phase instructions detailed above was played after task block 1, for task block 2 to 5.

**Table B.3**

*Sequences presented in the AGL task*

| 3-5 item grammatical sequences | 3-5 item ungrammatical sequences | 6-7 item grammatical sequences | 6-7 item ungrammatical sequences |
|---|---|---|---|
| 224 | 242 | 566534 | 256624 |
| 554 | 2432 | 233324 | 225664 |
| 5534 | 2423 | 553654 | 555364 |
| 5654 | 2342 | 566654 | 262534 |
| 2324 | 5435 | 232654 | 565634 |
| 22654 | 56354 | 226654 | 2332564 |
| 56534 | 25624 | 226534 | 2263354 |
| 23324 | 22564 | 2333324 | 2362534 |
| 56654 | 53564 | 2326534 | 2623354 |
| | 26254 | 5666534 | 2636524 |
| | 55364 | 5666654 | 2353264 |
| | | 5536654 | 2566234 |
| | | 5536534 | 5355664 |
| | | 2326654 | 5566354 |
| | | 2266534 | 2326354 |
| | | 2266654 | 2236564 |
| | | 5653654 | 2652364 |
| | | 2332654 | 5535664 |
| | | | 5536354 |

*Note.* Numbers were replaced by pictures of animals/color segments. Grammatical sequences followed AG rules (see Fig. 2) and were presented in learning and test phases. Ungrammatical sequences violated AG rules and were presented in test phases. Each sequence was displayed with an additional first element in the very beginning that completed the illustration of a train or a flag, respectively (see number 1 in Table B.1).

Supplementary Information: VISUAL ARTIFICIAL GRAMMAR LEARNING ACROSS ONE YEAR IN 7-YEAR-OLDS AND ADULTS

**Appendix C.**

**Control analyses without the first task block (task block 2-5)**

**Table C.1**

**A** *Year 1: Proportion Correct Across Block 2-5 in AGL per Session and Age Group*

| | **Session 1** | | **Session 2** | | **Session 3** | | **Transfer 1** | |
|---|---|---|---|---|---|---|---|---|
| | 7yo | Ad 1 | 7yo | Ad 1 | 7yo | Ad 1 | 7yo | Ad 1 |
| *N* | 27 | 28 | 27 | 28 | 27 | 28 | 27 | 28 |
| *M* | .56 | .82 | .62 | .91 | .63 | .94 | .64 | .90 |
| *SD* | .11 | .10 | .12 | .07 | .17 | .07 | .11 | .11 |
| Min | .23 | .63 | .40 | .78 | .37 | .76 | .33 | .65 |
| Max | .73 | 1.00 | .85 | 1.00 | .95 | 1.00 | .88 | 1.00 |
| *N** | 16 | 20 | 16 | 20 | 16 | 20 | 16 | 20 |
| *M* | .58 | .81 | .67 | .93 | .68 | .95 | .68 | .91 |
| *SD* | .08 | .11 | .12 | .06 | .18 | .07 | .09 | .12 |
| Min | .45 | .63 | .50 | .80 | .43 | .78 | .53 | .65 |
| Max | .73 | .98 | .85 | 1.00 | .95 | 1.00 | .88 | 1.00 |

**B** *Year 2: Proportion Correct in AGL Across Block 2-5 per Session and Age Group*

| | **Session 4** | | **Session 5** | | **Session 6** | | **Transfer 2** | |
|---|---|---|---|---|---|---|---|---|
| | 7yo | Ad 1 | 7yo | Ad 1 | 7yo | Ad 1 | 7yo | Ad 1 |
| *N* | 16 | 20 | 16 | 20 | 16 | 20 | 16 | 20 |
| *M* | .73 | .93 | .76 | .94 | .79 | .94 | .73 | .98 |
| *SD* | .11 | .10 | .17 | .10 | .15 | .09 | .14 | .09 |
| Min | .53 | .60 | .43 | .68 | .48 | .70 | .48 | .70 |
| Max | .93 | 1.00 | 1.00 | 1.00 | 1.00 | 1.00 | .98 | 1.00 |

*Note.* 7yo = 7-year-olds, Ad 1 = Adults, Min = minimal value, Max = maximal value.

* Subgroup of returning participants with data for Year 1 & Year 2.

**Table C.2**

*Results of rmANOVAs on Block 2-5 with Age (7-year-olds, Adults) as between subject factor & Session as within-subject factor*

| Session Comparison | Main Effects *Age* & *Session* | Interaction Effect *Age*Session* | Comparison to Result Pattern with all Task Blocks (Block 1-5) |
|---|---|---|---|
| Session 1 vs. Session 3 | both $F(1, 53) \geq 22.78$, both $p < .001$, both $\eta^2_g \geq .15$, both $BF_{incl} > 100$ | $F(1, 53) = 1.14$, $p = .290$, $\eta^2_g < .01$, $BF_{incl} = .41$ | Replicated |
| Session 1 vs. Transfer 1 | both $F(1, 53) \geq 17.06$, both $p < .001$, both $\eta^2_g \geq .12$, both $BF_{incl} > 100$ | $F(1, 53) < 0.01$, $p = .934$, $\eta^2_g < .01$, $BF_{incl} = .26$ | Replicated |
| Session 1 vs. Session 4 | both $F(1, 34) \geq 29.17$, both $p < .001$, both $\eta^2_g \geq .27$, both $BF_{incl} > 100$ | $F(1, 34) = .20$, $p = .532$, $\eta^2_g = .01$, $BF_{incl} = .29$ | Replicated |
| Session 3 vs. Session 4 | *Age* ($F(1, 34) = 41.52$, $p < .001$, $\eta^2_g = .51$, $BF_{incl} > 100$); *Session* ($F(1, 34) = 1.51$, $p = .227$, $\eta^2_g < .01$, $BF_{incl} = .31$) | $F(1, 34) = 6.18$, $p = .018$, $\eta^2_g = .03$, $BF_{incl} = 2.96$ | Replicated, but additional significant interaction *Age*Session* without block 1 (Post-hoc Test: 7-year-olds trend for Session 4 > Session 3: $t(15) = -1.89$, $p = .079$, $d = 0.47$, $BF_{10} = 1.07$; adults Session 3 vs. Session 4: $t(19) = 1.46$, $p = .160$, $d = 0.33$, $BF_{10} = .58$)) |

**Table C.3**

*Results of rmANOVAs on Block 2-5 with Age (7-year-olds, Adults) as between subject factor, Year & Session as within-subject factor*

| Session Comparison | Main Effects *Age, Year* & *Session* | Interaction Effects *Age*Year*Session, Age*Year, Age*Session, Year*Session* | Comparison to Result Pattern with all Task Blocks (Block 1-5) |
|---|---|---|---|
| Learning Gains Stimulus Set 1 (Year 1: Session 1 vs. Session 3, Year 2: Session 4 vs. Session 6) | all $F(1, 34) \geq 45.28$, all $p < .001$, all $\eta^2_g \geq .17$, all $BF_{incl} > 100$ | significant interactions *Year*Session* ($F(1, 34) = 12.35$, $p < .001$, $\eta^2_g = .04$, $BF_{incl} = 20.96$) & *Age*Year* ($F(1, 34) = 6.09$, $p = .019$, $\eta^2_g = .03$, $BF_{incl} = 5.63$); all other $F(1, 34) \leq 3.07$, $p \geq .089$, $\eta^2_g \leq .01$, $BF_{incl} \leq .85$ | Replicated |
| Transfer Savings Stimulus Set 2 (Year 1: Session 1 vs. Transfer 1, Year 2: Session 4 vs. Transfer 2) | all $F(1, 34) \geq 17.02$, all $p < .001$, all $\eta^2_g = .06$, all $BF_{incl} > 100$ | *Year*Session* $F(1, 34) = 16.99$, $p < .001$, $\eta^2_g = .06$, $BF_{incl} > 100$; all other $F(1, 34) \leq 1.43$, $p > .239$, $\eta^2_g \leq .06$, $BF_{incl} \leq .60$ | Replicated |

**Appendix D.**

**Performance correlations with language & memory skills, and with explicit knowledge**

**Table D.1**

*Year 1: Correlations for Memory Skills, Grammar Skills & Explicit Sequence Knowledge with AGL Performance*

| | **Learning Gains AGL** (Session 3 – Session 1) | | **Transfer Savings AGL** (Transfer 1 – Session 1) | | **Transfer Loss AGL** (Transfer 1 – Session 3) | |
|---|---|---|---|---|---|---|
| | 7-year-olds (*n* = 27) | Adults (*n* = 28) | 7-year-olds (*n* = 27) | Adults (*n* = 28) | 7-year-olds (*n* = 27) | Adults (*n* = 28) |
| **German Grammar I** (Plural) | .29 | -.27 | .03 | -.19 | ***-.32*** | .07 |
| **German Grammar II** (General) | .10 | -.06 [a] | .02 | -.14 [a] | -.06 [a] | -.22 |
| **Declarative Memory** | | | | | | |
| **Encoding** | -.16 | -.24 | .12 | -.23 | .22 | -.25 |
| **Retrieval** | -.03 | .09 | .02 | .01 | .06 | -.27 |
| **Working Memory** | .57* | .04 | ***.41*** | -.01 | -.21 | .01 |
| **Explicit knowledge** | ***.33*** | .10 | ***.42*** | .25 | .07 | .24 |

*Note.* AGL = Artificial Grammar Learning Task, ***italic =*** | r | ≥ .30, gray = $BF_{10}$ ≥ 3.

* corrected *p* < .05.

[a] The result pattern for German Grammar II with AGL performance remained the same when excluding *n* = 3 adults with very low scores in this subscale (*T* < 15).

**Table D.2**

*Correlations for Memory Skills, Grammar Skills (Year 1) & Explicit Sequence Knowledge with AGL Performance (Year 2)*

| | **Learning Gains AGL** (Session 6 – Session 4) | | **Transfer Savings AGL** (Transfer 2 – Session 4) | | **Transfer Loss AGL** (Transfer 2 – Session 6) | |
|---|---|---|---|---|---|---|
| | 7-year-olds (*n* = 16) | Adults (*n* = 20) | 7-year-olds (*n* = 16) | Adults (*n* = 20) | 7-year-olds (*n* = 16) | Adults (*n* = 20) |
| **German Grammar I** (Plural) | .22 | -.10 | ***.36*** | -.18 | .33 | -.13 |
| **German Grammar II** (General) | -.04 | -.28 | ***.46*** | -.11 | .36 | .29 |
| **Declarative Memory** | | | | | | |
| **Encoding** | .07 | .08 | .23 | -.08 | .29 | -.15 |
| **Retrieval** | ***.40*** | -.27 | ***.49*** | -.13 | .13 | .20 |
| **Working Memory** | ***.31*** | ***-.38*** | .32 | -.19 | .02 | .11 |
| **Explicit knowledge** | | | | | | |
| **Year 1** | -.22 | -.19 | .09 | ***-.38*** | .27 | ***-.30*** |
| **Year 2** | -.16 | -.14 | ***-.31*** | -.08 | -.12 | .14 |

*Note.* AGL = Artificial Grammar Learning Task, ***italic*** **=** | r | ≥ .30.
